# Nephrotoxicity of the BRAF-kinase inhibitor Vemurafenib is driven by off-target Ferrochelatase inhibition

**DOI:** 10.1101/2021.01.29.428783

**Authors:** Yuntao Bai, Ji Young Kim, Laura A. Jayne, Megha Gandhi, Kevin M. Huang, Josie A. Silvaroli, Veronika Sander, Jason Prosek, Kenar D. Jhaveri, Sharyn D. Baker, Alex Sparreboom, Amandeep Bajwa, Navjot Singh Pabla

**Author notes:** ***Correspondence should be addressed to:*** Navjot Pabla, Division of Pharmaceutics and Pharmacology, College of Pharmacy and Cancer Center, 460 W 12th Ave, Columbus, OH 43221, USA.

## Abstract

A multitude of disease and therapy related factors drive the frequent development of renal disorders in cancer patients. Along with chemotherapy, the newer targeted therapeutics can also cause renal dysfunction through on and off-target mechanisms. Interestingly, among the small-molecule inhibitors approved for the treatment of cancers that harbor BRAF-kinase activating mutations, vemurafenib can trigger tubular damage and acute kidney injury (AKI). To investigate the underlying mechanisms, here, we have developed cell culture and mouse models of vemurafenib nephrotoxicity. Our studies show that at clinically relevant concentrations vemurafenib induces cell-death in transformed and primary murine and human renal tubular epithelial cells (RTEC). In mice, two weeks of daily vemurafenib treatment causes moderate AKI with histopathological characteristics of RTEC injury. Importantly, RTEC-specific BRAF gene deletion did not influence renal function under normal conditions or alter the severity of vemurafenib-associated renal impairment. Instead, we found that inhibition of ferrochelatase (FECH), an enzyme involved in heme biosynthesis contributes to vemurafenib nephrotoxicity. FECH overexpression protected RTECs and conversely FECH knockdown increased the sensitivity to vemurafenib nephrotoxicity. Collectively, these studies suggest that vemurafenib-associated RTEC dysfunction and nephrotoxicity is BRAF-independent and caused in part by off-target FECH inhibition.

**Translational Statement:** BRAF is the most frequently mutated protein kinase and a critical oncogenic driver in human cancers. In melanoma and other cancers with BRAF activating mutations, BRAF targeted small-molecule therapeutics such as vemurafenib, and dabrafenib have shown remarkable clinical benefits. However, recent clinical studies have shown that a significant number of patients that receive vemurafenib develop AKI through mechanisms that remain unknown. The present study describes the development of novel experimental models of vemurafenib nephrotoxicity and reveals the underlying off-target mechanisms that contribute to renal injury.

## Introduction

A broad array of factors increase the risk of renal dysfunction in cancer patients^1–3^. While certain malignancies can directly affect kidney function^4^, cancer therapy can also trigger fluid and electrolyte disorders^5^, acute kidney injury (AKI)^6^, and chronic kidney disease (CKD)^7^. Renal disorders jeopardize the continuation of cancer therapy and contribute to the development of debilitating short and long-term adverse sequelae. Along with chemotherapeutics, targeted antibodies^8^ and small-molecule inhibitors^9^ are also associated with renal impairment. Thus, one of the challenges in the emerging field of onconephrology^10–13^ is to determine the mechanisms associated with these toxicities in order to develop mitigating strategies.

Dysregulation of the mitogen-activated protein kinase (MAPK) pathway is a major driver of oncogenesis^14^. The Raf (rapidly accelerated fibrosarcoma) family of serine/threonine kinases act as a conduit between upstream Ras signaling and downstream MAPK kinase activation, relaying signaling cues from the extracellular environment and directing cell proliferation, differentiation, migration and survival^15^. Mammals possess three RAF proteins: RAF1 (CRAF), ARAF and BRAF, which play essential and distinct physiological roles. Importantly, somatic activating mutations in BRAF are frequent in hairy-cell leukemia (100%), melanoma (50%– 60%), and thyroid cancer (40%– 60%)^15,16^. The recent development of orally bioavailable BRAF-inhibitors, namely vemurafenib^17,18^ and dabrafenib^19^ have brought exceptional clinical benefits, especially in melanoma patients.

Dermatological toxicities are the major side effects related to vemurafenib treatment^20^. While renal toxicities were not observed in the clinical trials^21^, a growing body of literature suggests that a significant percentage of patients treated with vemurafenib can develop AKI^22–24,24–26^. Acute tubular necrosis, electrolyte disorders, and subnephrotic-range proteinuria have been reported in a subset of vemurafenib-treated patients^22,25^. These studies have shown that patients treated with vemurafenib as compared to dabrafenib have a higher incidence of AKI and analysis of kidney biopsies have established tubular injury as the major histopathological lesion^22^.

Due to paucity of experimental models, the mechanisms underlying vemurafenib nephrotoxicity remain unclear. Furthermore, it is unknown if BRAF is essential for renal tubular function. It is also unknown whether vemurafenib nephrotoxicity is BRAF-dependent. To address these key questions, here we have developed a renal tubule-specific BRAF knockout mice and established *in vitro* and *in vivo* models of vemurafenib nephrotoxicity. Our studies suggest that BRAF knockout in renal tubules does not trigger renal impairment and vemurafenib-associated RTEC dysfunction and AKI is BRAF-independent.

## Results

### Vemurafenib induces cell-death in cultured RTECs

We initially sought to determine if vemurafenib could cause direct toxicity in cultured RTECs. To this end, we treated transformed tubular epithelial cells of murine (BUMPT) and human origin (HK-2) with vemurafenib and tested its effect on cellular survival. In these experiments, we also included cisplatin, a well-studied nephrotoxic drug^27^, as well as BRAF inhibitor debrafenib, and the multi-kinase and CRAF-inhibitor sorafenib^28^. Similar to cisplatin, vemurafenib treatment reduced cellular viability as measured by trypan blue staining (**Fig. 1A-B**) in both BUMPT and HK-2 cells (IC_50_=~50μM). Notably, pharmacokinetic studies in humans have shown that plasma levels of vemurafenib can range from 50-100 μM^21^. Confirmatory experiments with MTT assays showed that at 50 μM concentration, vemurafenib can cause ~50% reduction in cellular viability within 48 hours (**Suppl. Fig.1A-B**), with a parallel increase in caspase activation (**Fig. 1C-D**).

**Figure 1:**
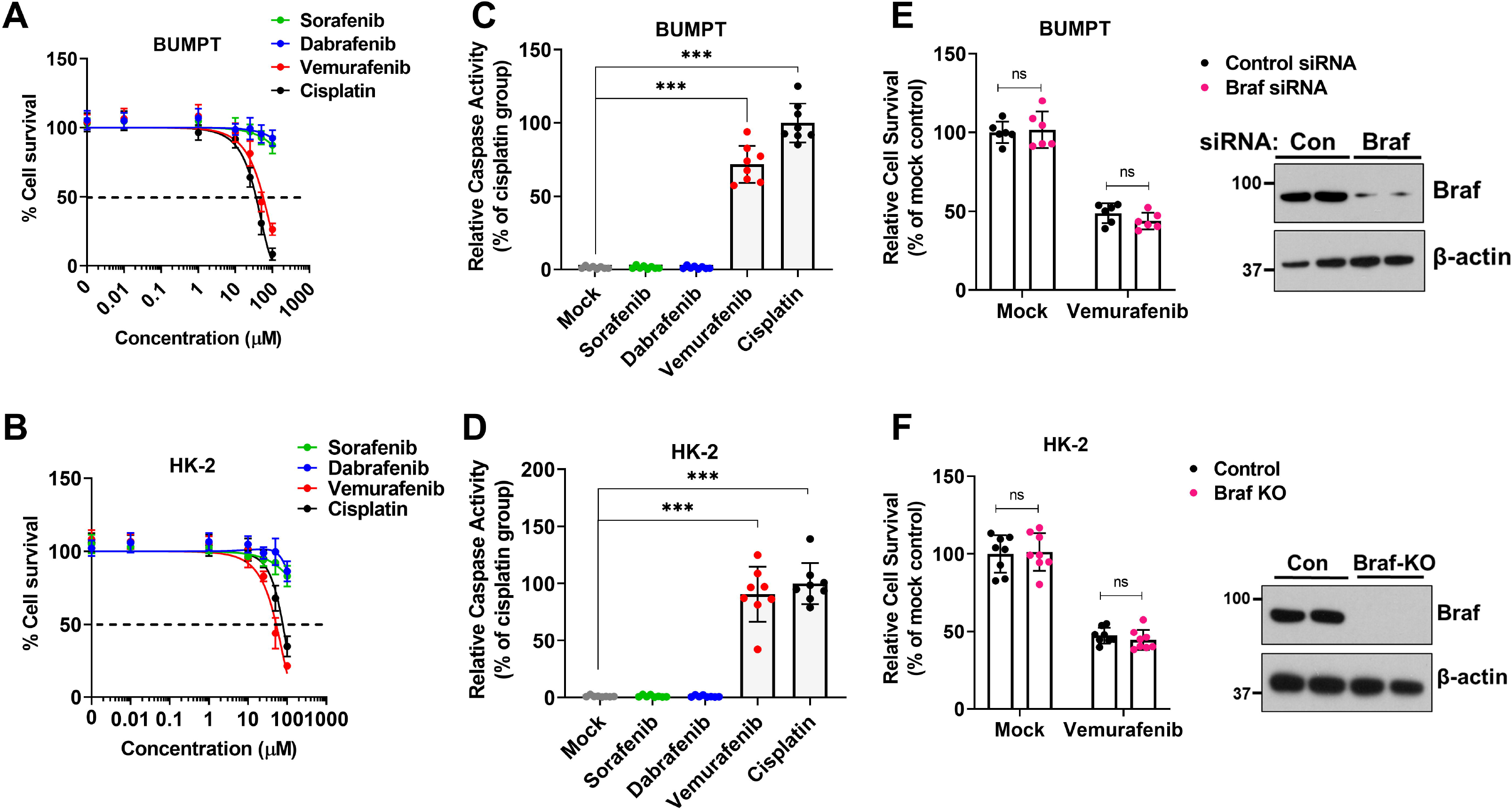
Vemurafenib induces cell death in murine and human tubular epithelial cell lines. Tubular epithelial cell lines of murine (BUMPT) and human (HK-2) origin were treated with vehicle, cisplatin or kinase inhibitors including vemurafenib followed by assessment of cell viability and cell death. (**A-B**) Dose response experiments (0-100 μM) and trypan blue based cellular viability assays at 48 hours post treatment showed that veumurafenib can induce cell death in RTEC cell lines with IC50 values of approximately 50 μM. Survival data was normalized to vehicle group and are presented as mean (n□=□5 biologically independent samples), from one out of three independent experiments, all producing similar results. (**C-D**) BUMPT and HK-2 cells were treated with vehicle or indicated drugs at 50 μM concentration, followed by measurement of caspase activity at 48 hours. The results show that similar to cisplatin, vemurafenib can reduce RTEC viability. Data are presented as individual data points (n_=_8 biologically independent samples from three independent experiments). (**E**) RNAi mediated Braf knockdown in BUMPT cells did not influence vemurafenib associated cell death (50 μM for 48 hours) as assessed by trypan blue based viability assay. Data are presented as individual data points (n□=□8 biologically independent samples from three independent experiments). A representative immunoblot (right panel) shows the successful knockdown of Braf gene. (**F**) CRISPR/Cas9 mediated Braf knockout in HK-2 cells did not influence vemurafenib associated cell death (50 μM for 48 hours) as assessed by trypan blue based viability assay. Data are presented as individual data points (n□=□8 biologically independent samples from three independent experiments). A representative immunoblot (right panel) shows the successful knockout of Braf gene. In all the graphs (n=5-8 biologically independent samples), experimental values are presented as mean□±□s.d. The height of error bar□=□1 s.d. and p□<□0.05 was indicated as statistically significant. One-way ANOVA followed by Dunnett’s was carried out and statistical significance is indicated by *p < 0.05, **p < 0.01, ***p < 0.001.

Intriguingly, kinase assays showed that at 50 μM concentration vemurafenib, debrafenib, and sorafenib inhibited BRAF-kinase activity to similar levels (**Suppl. Fig.1C-D**), however, under these conditions only vemurafenib triggered RTEC cell-death. We next questioned whether BRAF kinase is essential for RTEC survival and a causal factor in vemurafenib-induced cell-death. To address this, we carried out RNAi-mediated knockdown and CRISPR/Cas9-mediated BRAF knockout in BUMPT and HK-2 cells respectively (**Fig. 1E-F**). BRAF knockdown or knockout had no impact on cellular viability under normal conditions and neither did it influence vemurafenib-induced cell-death (**Fig. 1E-F and Suppl. Fig. 2**). These results suggest that vemurafenib can trigger cell-death in cultured RTECs, seemingly through a BRAF-independent mechanism.

**Figure 2:**
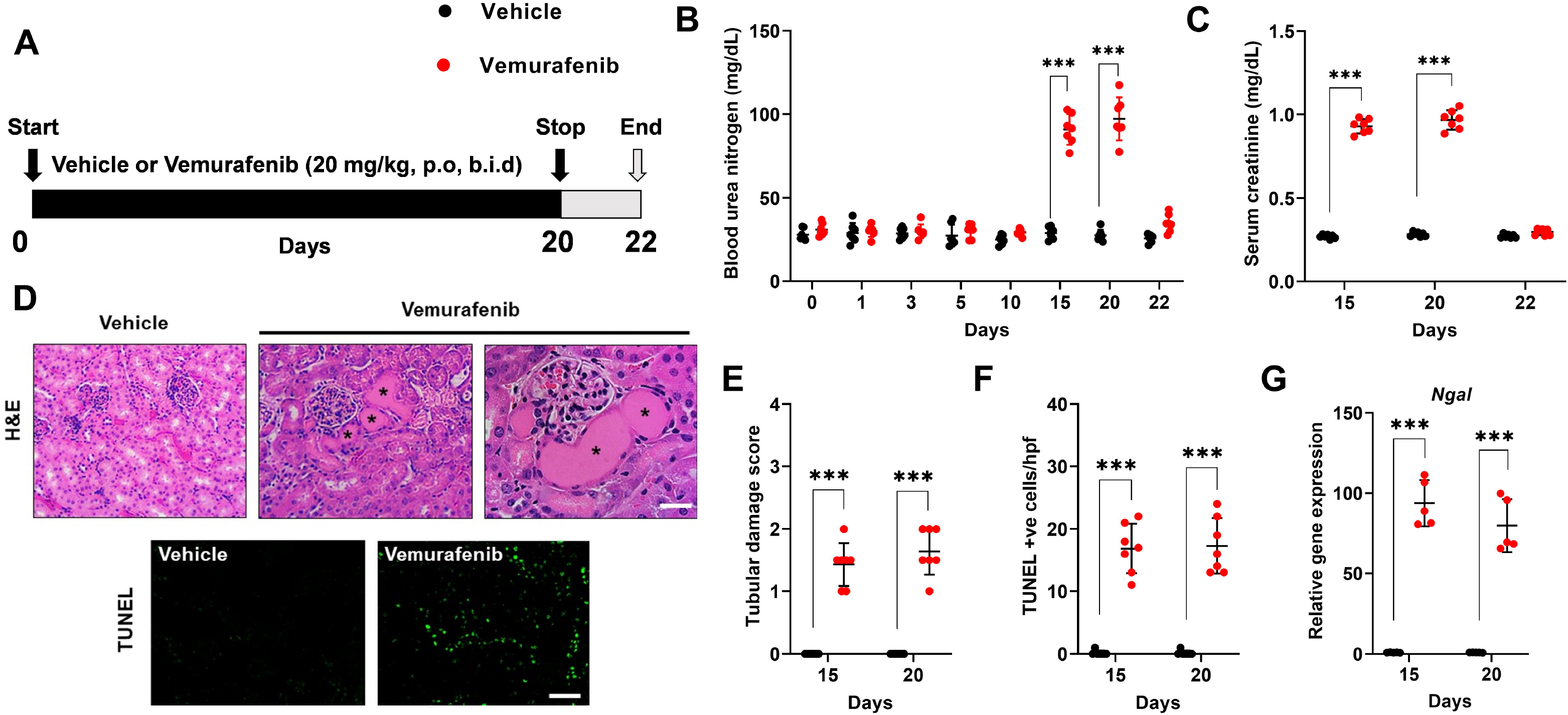
Development of a mouse model of vemurafenib nephrotoxicity. (**A**) Schematic representation of dosing strategy. Age-matched, 8-12 weeks old male C57BL/6J mice were treated with either vehicle or 20 mg/kg vemurafenib (p.o, b.i.d.) for 20 days followed by cessation of drug administration for 2 days and subsequent endpoint analysis of renal function. (**A**) Analysis of blood urea nitrogen levels showed that vemurafenib can induce AKI after 2 weeks of continuous treatment and cessation of drug adminstration reversed the increase in BUN levels. (**C**) Vemurafenib treatment also resulted in increased serum creatinine levels indicating significant renal impairment (**D-E**) Histological analysis of renal tissues showed that vemurafenib treated mice had clear tubular epithelial injury and cell death. Representative H&E staining depicting renal tubular damage (indicated by an asterisk) linked with vemurafenib-associated AKI. (**D & F**) TUNEL staining of renal tissues revealed significant tubular epithelial cell death in the vemurafenib treated mice. (**G**) Renal NGAL gene expression analysis further confirmed significant renal damage in vemurafenib treated mice. In all the bar graphs (n=7 biologically independent samples), experimental values are presented as mean□±□s.d. The height of error bar□=□1 s.d. and p□<□0.05 was indicated as statistically significant. Student’s t-test was carried out and statistical significance is indicated by *p < 0.05, **p < 0.01, ***p < 0.001. Scale bar (D): 100□μm.

### Establishment of a mouse model of vemurafenib nephrotoxicity

To establish a murine model of vemurafenib nephrotoxicity, we initially performed a pharmacokinetic study in C57B6/J mice. In concordance with previous work^29^, we found that at 20 mg/kg dose, plasma drug (~50μM) concentrations (**Suppl. Fig. 3**) were similar to those observed in humans^21^. We then determined the effect of 20 mg/kg twice-daily vemurafenib administration on kidney function (**Fig. 2A**). We noticed a significant increase in blood urea nitrogen (**Fig. 2B**) and serum creatinine (**Fig. 2C**) levels after 2 weeks of b.i.d. dosing. Histological examination (**Fig. 2D-E**) revealed tubular necrosis and TUNEL staining (**Fig. 2 D&F**) showed significant tubular epithelial cell-death. In C57B6/J mice, several injury^30^, repair^31^ and inflammatory genes^32–35^ are upregulated during ischemia, cisplatin, and rhabdomyolysis associated AKI^36^. We found a similar increase in injury, repair, and inflammation-related genes during vemurafenib nephrotoxicity (**Fig. 2G and Suppl. Fig. 4**). Similar to humans^22,25^, we found that treatment discontinuation in mice reverses vemurafenib-associated renal impairment (day 20 versus 22).

### Vemurafenib nephrotoxicity is BRAF-independent

To evaluate the RTEC-specific role of BRAF kinase, we generated conditional knockout mice (BRAF^PT-/-^) by crossing the BRAF floxed^37^ mice with the Ggt1-Cre mice^38^. We found that BRAF deficiency (**Figure 3A**) did not influence normal renal function (**Suppl. Fig. 5**). Moreover, when the control and BRAF^PT-/-^ littermates were challenged with vemurafenib, the extent of renal damage as evaluated by blood urea nitrogen, serum creatinine, as well as histological damage and injury biomarker analysis was found to be similar in both groups (**Figure 3B-F**). To corroborate the *in vivo* findings, we isolated primary RTECs from BRAF floxed mice and carried out in vitro Cre-mediated deletion (**Suppl. Fig. 6A**). Vemurafenib treatment induced cell-death in primary RTECs and BRAF gene-deletion did not influence cell-death under these conditions (**Suppl. Fig. 6B**). These findings support the notion that BRAF-kinase inhibition is unlikely to be the underlying cause of vemurafenib-associated AKI.

**Figure 3:**
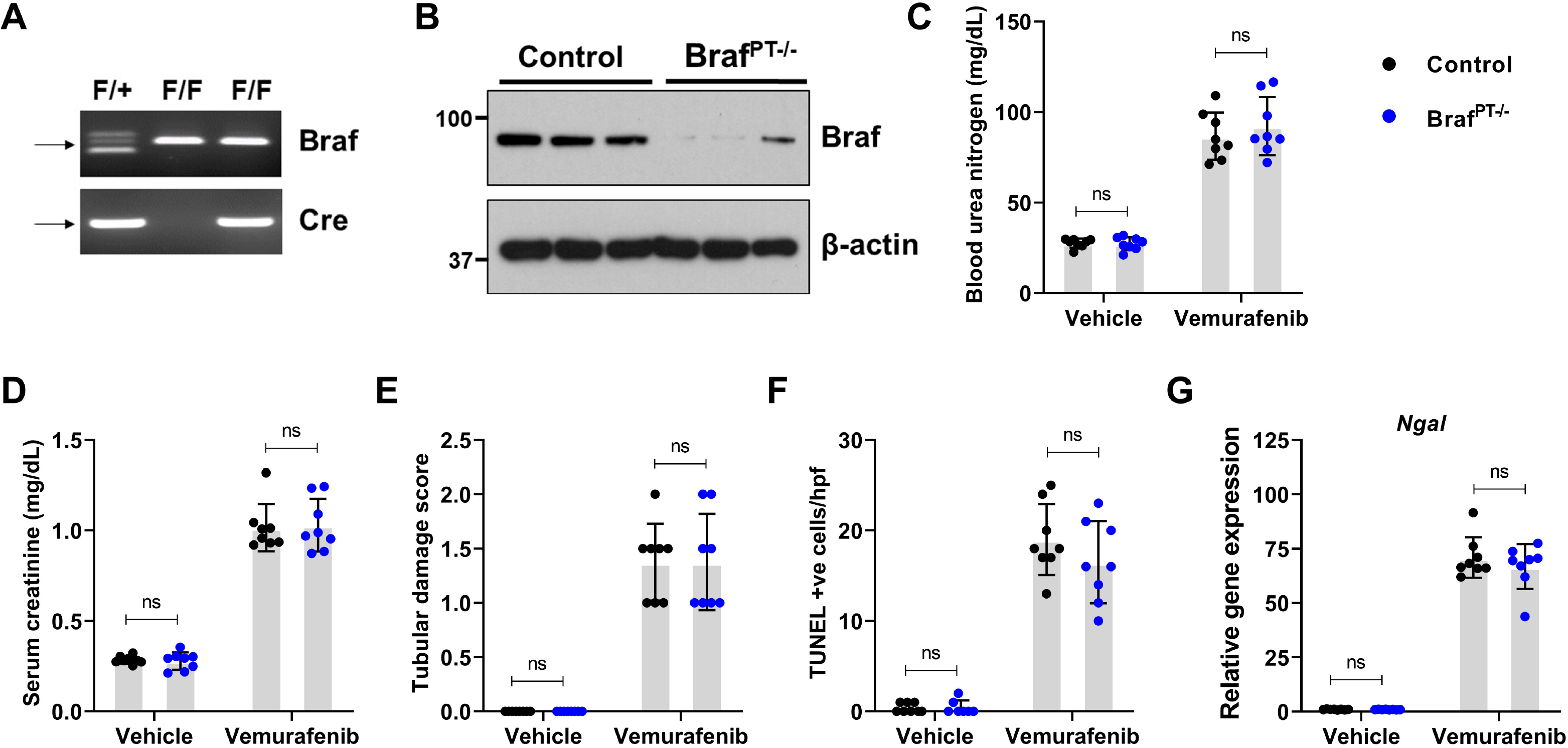
Vemurafenib nephrotoxicity is not influenced by RTEC-specific Braf gene deletion. To generate mice with renal tubule-specific *Braf* knockout, *Ggt1-Cre* mice were crossed with *Braf-*floxed mice. **(A-B)** Representative genotyping and immunoblots showing successful knockout in the renal tissues. Littermate control and *Braf* conditional knockout mice (indicated by Braf^PT−/−^) were used to study the role of Braf in vemurafenib nephrotoxicity. Age-matched, 8-12 weeks old male littermate mice were treated with either vehicle or 20 mg/kg vemurafenib (p.o, b.i.d.) for 20 days followed by subsequent endpoint analysis of renal function. Blood urea nitrogen (**C**), serum creatinine (**D**), and histological analysis (**E**) were performed to examine renal function and damage. The extent of functional renal impairment and damage was similar between the control and Braf deficient mice. (**F**) TUNEL staining also showed similar amount of renal epithelial cell death in the control and knockout tissues. (**G**) Renal NGAL gene expression analysis. In all the bar graphs (n=8 biologically independent samples), experimental values are presented as mean□±□s.d. The height of error bar□=□1 s.d. and p□<□0.05 was indicated as statistically significant. One-way ANOVA followed by Tukey’s multiple-comparison test was carried out and statistical significance is indicated by *p < 0.05, **p < 0.01, ***p < 0.001.

### Identification of molecular targets associated with vemurafenib toxicity

Small-molecule kinase inhibitors can target the ATP-binding pocket; however, the high conservation of the ATP binding site within the kinase families poses a significant challenge in developing highly specific kinase inhibitors^39^. Previous studies^40,41^ have systematically profiled the promiscuity of kinase inhibitors including vermurafenib. To ascertain if inhibition of these kinases contribute to vermurafenib cytotoxicity in RTECs, we used a chemical genetics approach^42^. BUMPT cells were transfected with plasmids encoding wild-type or inhibitor-resistant kinase genes and their effect on cellular survival was monitored (**Suppl. Fig. 7**). Surprisingly, inhibitor-resistant kinase overexpression did not rescue cytotoxicity in BUMPT cells (**Suppl. Fig. 7**), raising the possible role of a non-kinase related mechanism.

Interestingly, chemical proteomics studies^43,44^ have shown that vemurafenib but not dabrafenib can inhibit ferrochelatase (FECH). FECH is a mitochondrial protein that catalyzes the insertion of ferrous iron into protoporphyrin IX, and is the terminal enzyme involved in heme biosynthesis^45^. To ascertain if FECH inhibition causes vemurafenib-associated RTEC cell-death, we initially quantified FECH activity in BUMPT and HK-2 cells. We found that vemurafenib significantly inhibited FECH activity in RTEC cell-lines (**Fig. 4A-B**). Importantly, wild-type FECH overexpression protected BUMPT and HK-2 from vemurafenib associated cell-death (**Fig. 4C-G**). Overexpression of an inactive FECH mutant (FECH^mut^) did not influence vemurafenib associated cell-death. These results indicate that FECH inhibition might contribute to RTEC cell-death under in vitro conditions.

**Figure 4:**
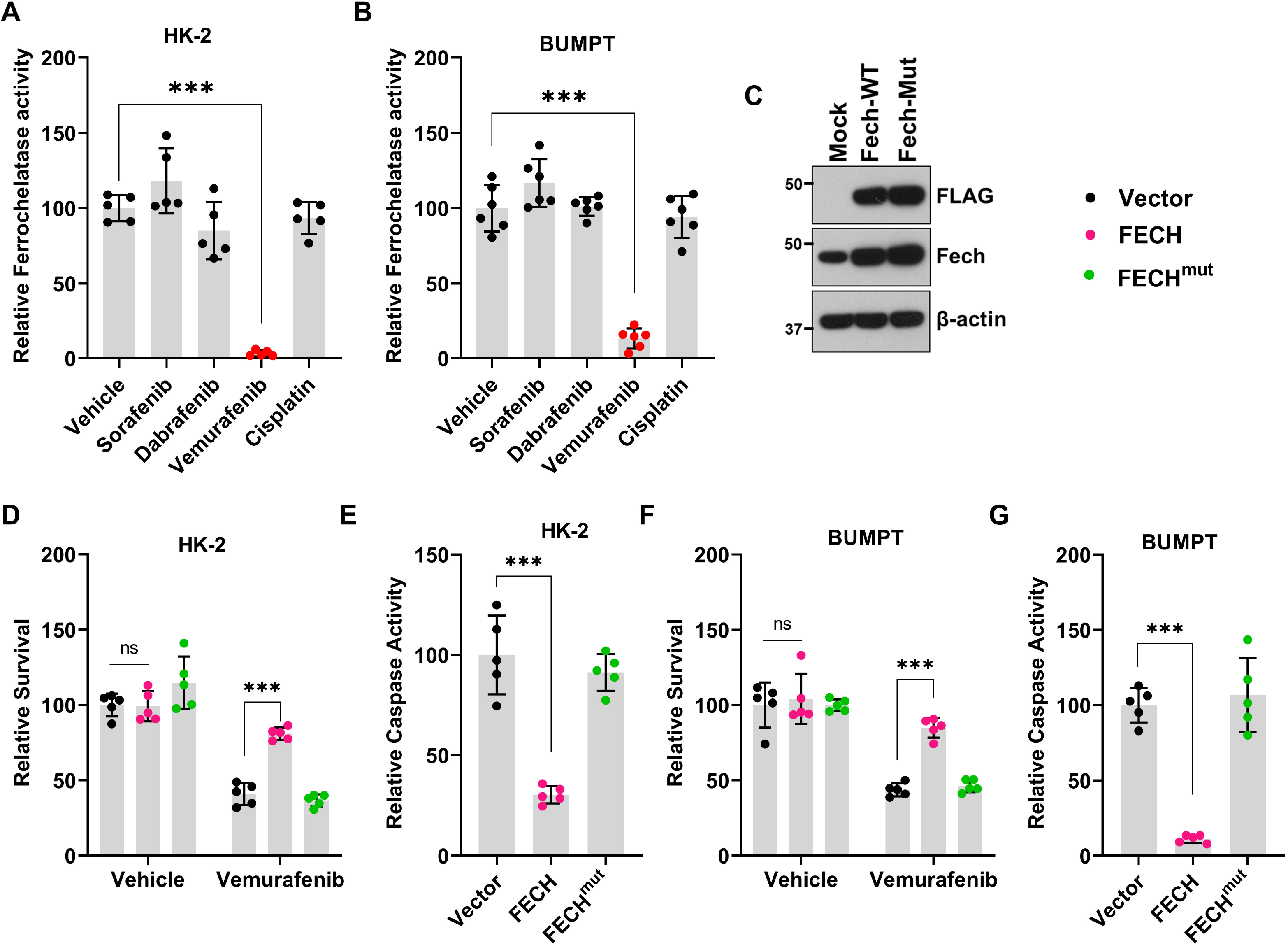
Ferrochelatase inhibition contributes to Vemurafenib mediated RTEC cell death. Tubular epithelial cell lines of murine (BUMPT) and human (HK-2) origin were treated with vehicle, cisplatin or kinase inhibitors including vemurafenib at 50 μM concentration, followed by assessment of ferrochelatase activity at 24 hours. (**A-B**) Ferrochelatase activity was inhibited by vemurafenib in BUMPT and HK-2 cells. (**B**) Representative immunoblot showing overexpression of FLAG-tagged wild type and mutant FECH. Blots are representative of three independent experiments. (**D-G**) Empty vector, wild type FECH, or FECH mutant (M98K) was overexpressed in BUMPT (transient transfection) and HK-2 (lentiviral transduction) cells followed by treatment with either vehicle or 50 μM vemurafenib for 48 hours. Trypan blue based survival assays and caspase assays showed that wild type FECH overexpression can protect BUMPT and HK-2 cells from vemurafenib-associated cell death. In all the bar graphs (n=5-6 biologically independent samples) from one out of three independent experiments, all producing similar results, and experimental values are presented as mean□±□s.d. The height of error bar□=□1 s.d. and p□<□0.05 was indicated as statistically significant. One-way ANOVA followed by Tukey’s multiple-comparison test was carried out and statistical significance is indicated by *p < 0.05, **p < 0.01, ***p < 0.001.

### In vivo FECH knockdown hastens vemurafenib nephrotoxicity

Immunofluorescence studies showed that FECH is highly expressed in the cortical tubular epithelial cells (**Suppl. Fig. 8A**). While FECH protein levels remained unaltered, enzymatic assays showed a progressive decline in FECH activity during vemurafenib nephrotoxicity (**Suppl. Fig. 8B-C**). Reduction in FECH activity was vemurafenib-specific and not a generalized consequence of AKI (**Suppl. Fig. 8D**). Given that a progressive decline in FECH activity preceded the development of AKI at 2 weeks, we questioned if FECH knockdown would influence vemurafenib nephrotoxicity. Using hydrodynamic siRNA injection approach^46,47^, we identified a specific siRNA that reduced FECH protein expression by ~70% (**Fig. 5A-B**). FECH knockdown did not influence normal renal function, however strikingly, mice with FECH knockdown developed vemurafenib nephrotoxicity within three days in contrast to 2 weeks in control mice (**Fig. 5C-F**). This increased sensitivity was vemurafenib-specific because no difference in the extent of renal impairment was observed when the mice were challenged with cisplatin (**Suppl. Fig. 9**). These results suggest that FECH knockdown can remarkably hasten the development of vemurafenib-associated AKI.

**Figure 5:**
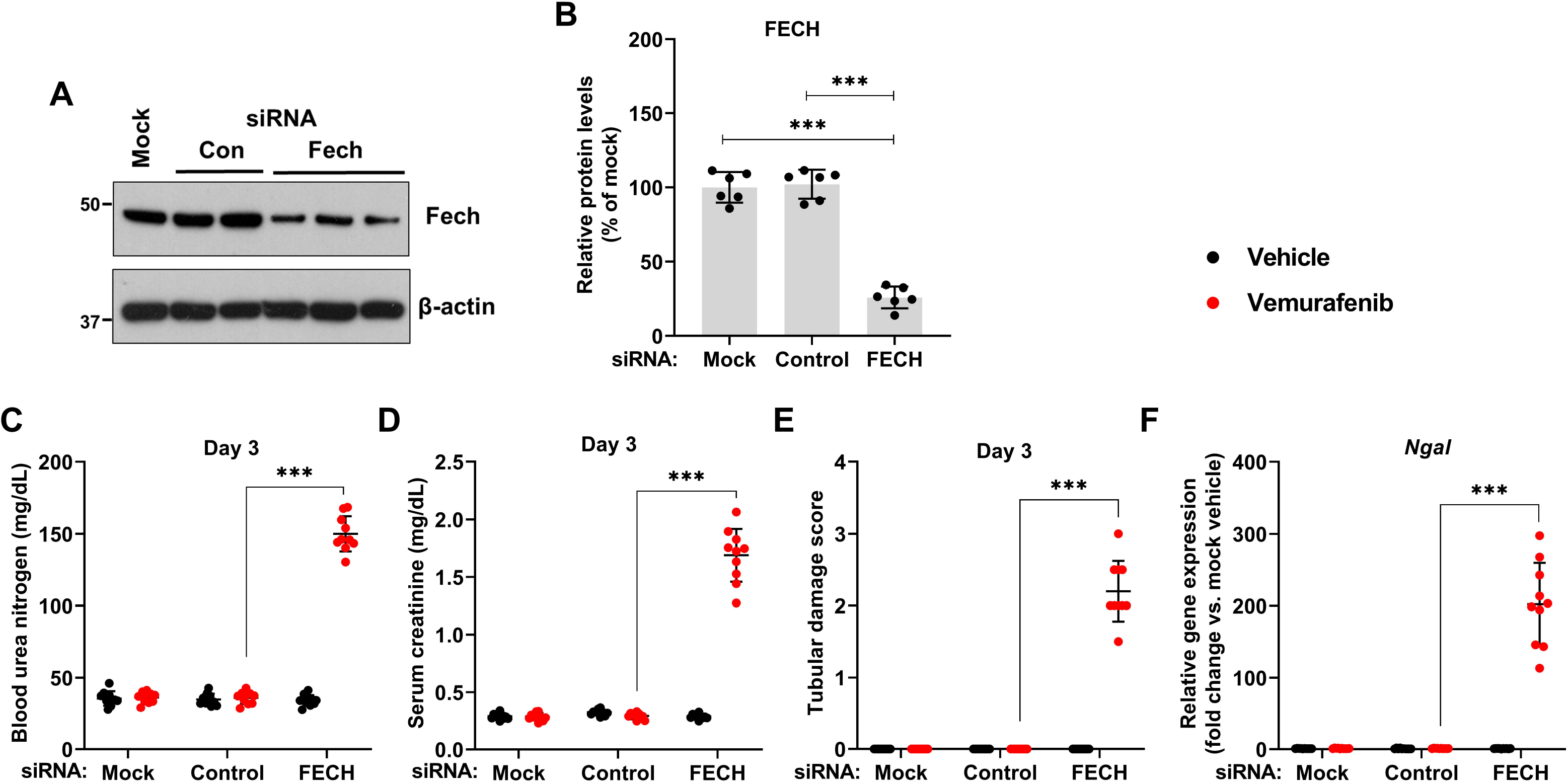
In vivo siRNA mediated FECH knockdown hastens the development of vemurafenib nephrotoxicity. Age-matched male (8-12 weeks) C57BL/6 mice were administered with three once-daily intravenous injections of control (non-specific) or FECH targeting siRNAs (25 μg in 0.5 ml of PBS). In one group (mock) 0.5 ml of PBS was injected. One day later mice were treated with either vehicle or 20 mg/kg vemurafenib (p.o, b.i.d.) for 3 days followed by endpoint analysis of renal function. (**A**) Representative blots and (**B**) densitometric analysis show that the targeted siRNA was able to knock-down FECH proteins levels by approximately 75%. Blots are representative of three independent experiments, all producing similar results. Blood urea nitrogen (**C**), serum creatinine (**D**), and histological analysis (**E**) Renal NGAL gene expression analysis (**F**) showed that the FECH knockdown mice developed vemurafenib nephrotoxicity within 3 days of treatment, while the control group demonstrated no obvious renal injury or damage. In all the bar graphs (n=6-10 biologically independent samples), experimental values are presented as mean□±□s.d. The height of error bar□=□1 s.d. and p□<□0.05 was indicated as statistically significant. One-way ANOVA followed by Tukey’s multiple-comparison test was carried out and statistical significance is indicated by *p < 0.05, **p < 0.01, ***p < 0.001.

### Accelerated development of vemurafenib nephrotoxicity in FECH mutant mice

In humans, mutations associated with reduced ferrochelatase activity can cause erythropoietic protoporphyria (EPP), a disease characterized by cutaneous photosensitivity and liver damage. In the mice with homozygous (fch/fch) mutations that reduce the FECH activity to 2.7-6.3%, photosensitivity and hepatic dysfunction is observed^45^. On the other hand, the heterozygotes (+/fch) mice have 45-65% of normal FECH activity and do not display any skin or liver damage. Congruent with a previous study^45^, we found that the heterozygous mutant mice had approximately 50% reduction in renal FECH activity (**Fig. 6A**). Next, we studied the consequence of reduced FECH activity on the development of vemurafenib nephrotoxicity. In comparison to control littermates, the heterozygous mice developed vemurafenib nephrotoxicity in an accelerated manner. At 7 days of vemurafenib treatment, when the control mice had no apparent renal impairment, BUN, serum creatinine, histological, and gene expression analysis (**Fig. 6 B-E**) showed a precipitous decline in renal function in the heterozygous mice. Given that vemurafenib inhibits FECH activity in RTECs and genetic reduction of FECH activity accelerates the development of AKI, we propose that inhibition of renal FECH function contributes to vemurafenib nephrotoxicity.

**Figure 6:**
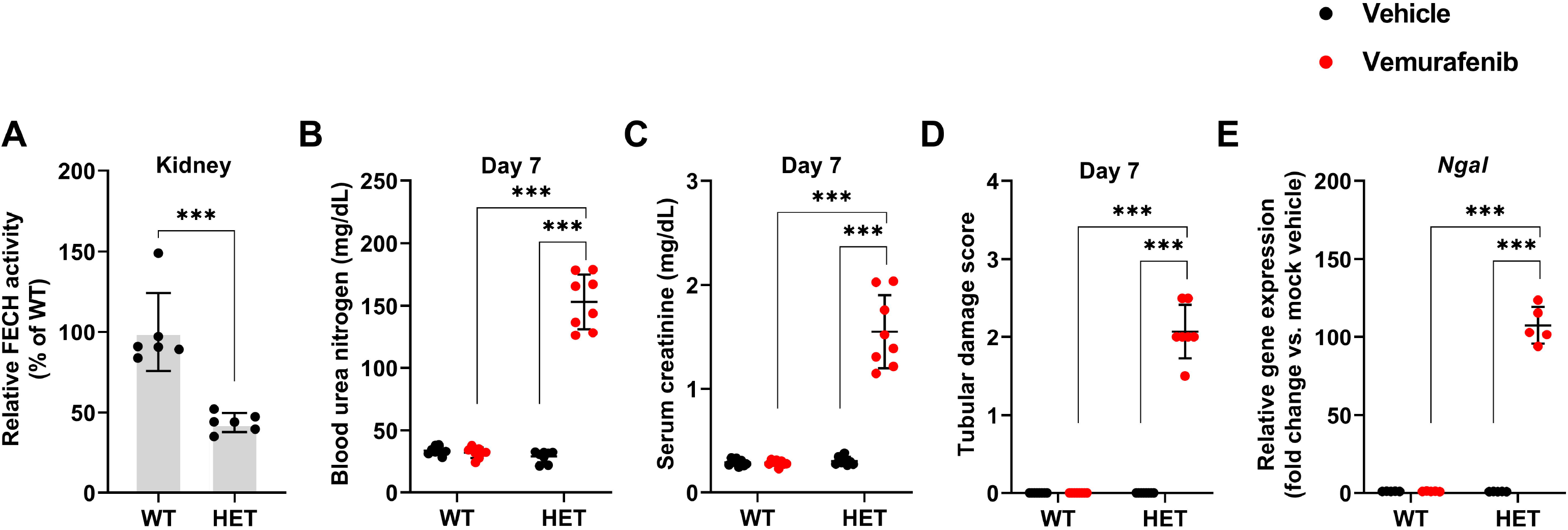
Accelerated development of vemurafenib nephrotoxicity in FECH mutant mice. (**A**) Renal tissues from wild type and heterozygous (Fch/+) mutant mice (littermates) were used to evaluate FECH activity using an ezymmatic assay. The result show an approximately 50% reduction in FECH activity in the heterozygous mice. The wild type and heterozygous mice were then challenged with either vehicle or 20 mg/kg vemurafenib (p.o, b.i.d.) for 7 days followed by endpoint analysis of renal function. Blood urea nitrogen (**B**), serum creatinine (**C**), and histological analysis (**D**) Renal NGAL gene expression analysis (**E**) showed that the FECH heterozygous mutant mice developed vemurafenib nephrotoxicity within 7 days of treatment, at a time-point when the wild type group demonstrated no obvious renal injury or damage. In all the bar graphs (n=6-8 biologically independent samples), experimental values are presented as mean_±_s.d. The height of error bar□=□1 s.d. and p□<□0.05 was indicated as statistically significant. One-way ANOVA followed by Tukey’s multiple-comparison test was carried out and statistical significance is indicated by *p < 0.05, **p < 0.01, ***p < 0.001.

### CRISPR/Cas9 mediated FECH knockout triggers cell-death in HK-2 cells

We questioned if genetic FECH inhibition would be sufficient to trigger cell-death in cultured renal tubular epithelial cells. To address this, we utilized a doxycycline-inducible CRISPR/Cas9 system, wherein we generated stable HK-2 cells that express either a control or FECH-targeted sgRNA, along with doxycycline-inducible Cas9. As shown in **Fig. 7A-B**, doxycycline treatment resulted in decline in FECH protein expression in the stable cells expressing the FECH targeted sgRNA. When we further monitored these cells for 24-72 hours post doxycycline induction, we noticed a clear decline in cellular viability in the FECH knockout cells as measured by trypan blue staining (**Fig. 7C**). At 72 hours, MTT and caspase activity assays confirmed reduced viability and increased cell-death in the FECH knockout cells (**Fig. 7D-E**). The observation that genetic FECH deficiency impairs RTEC viability supports the hypothesis that FECH inhibition could trigger RTEC cell-death and dysfunction.

**Figure 7:**
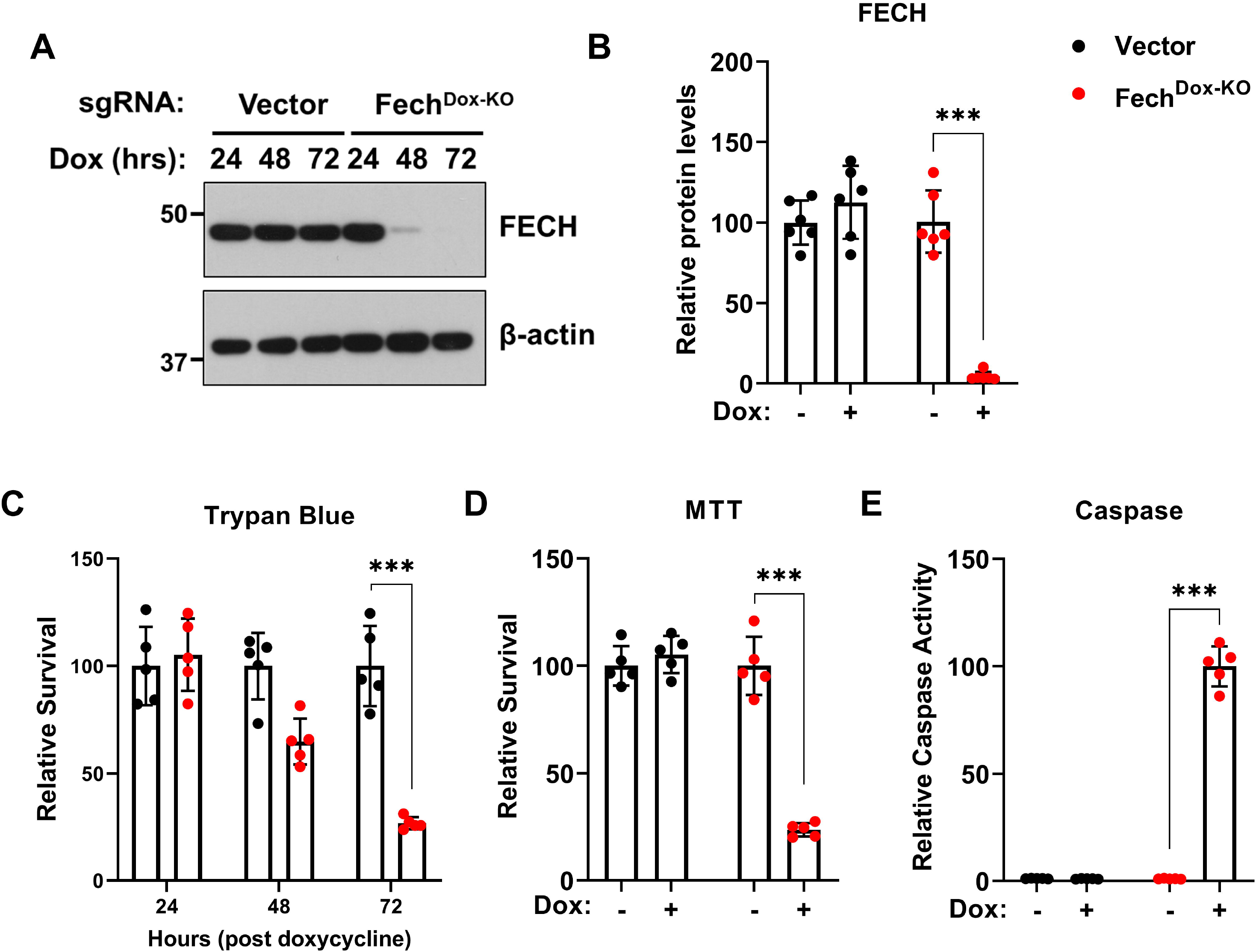
CRISPR/Cas9 mediated FECH knockout induces cell death in HK-2 cells. Stable HK-2 cells were generated with either an control or FECH targeted sgRNA. In these stable cells lines, Cas9 expression can be induced by doxycycline treatment. (**A**) In the presence of doxycycline, FECH gene deletion occurs in the cells that express FECH targed sgRNA. Representative immunoblot depicting successful gene knockout. Blots are representative of three independent experiments. (**B**) Densitometric analysis of FECH protein after 72 hours of doxycycline treatment. (**C**) Control (vector) and FECH sgRNA (FECH^Dox-KO^) stable cells growing in normal media, were treated with doxycycline and trypan blue based survival assays were performed at 24-72 hours. The results show that FECH knockdown resulted in significant reduction in cell survival. (In all the bar graphs (n=5-6 biologically independent samples) from one out of three independent experiments, all producing similar results, and experimental values are presented as mean□±□s.d. The height of error bar□=□1 s.d. and p□<□0.05 was indicated as statistically significant. One-way ANOVA followed by Dunnett’s test was carried out and statistical significance is indicated by *p < 0.05, **p < 0.01, ***p < 0.001.

## Discussion

In the current study, we have established cell culture and murine models of vemurafenib nephrotoxicity that have provided important mechanistic insights. Our studies reveal that vemurafenib triggers RTEC dysfunction and nephrotoxicity through a BRAF-independent and ferrochelatase-dependent manner.

Similar to most protein kinase inhibitors, vemurafenib is mainly excreted via feces (94%), with urinary excretion (1%) playing an insignificant role^21,48^. Based on our *in vitro* and *in vivo* studies, we propose that vemurafenib causes direct toxicity to tubular epithelial cells. Several lines of evidence suggest that this toxicity is independent of BRAF-kinase inhibition. Firstly, both vemurafenib and dabrafenib treatment resulted in BRAF inhibition, however only vemurafenib triggered RTEC cell-death. Secondly, BRAF deletion in murine and human RTECs did not influence cell survival under normal conditions. BRAF deletion or knockdown also did not influence vemurafenib-induced cell-death and AKI. This also eliminates the possibility that paradoxical MAPK activation initiated by inhibitor-induced wild type BRAF dimerization^49^ might contribute to nephrotoxicity. Thirdly, chemical genetic studies with transfection of vemurafenib-resistant BRAF gatekeeper mutants did not confer protection from vemurafenib-induced cell-death. While genetic compensation remains a possibility, collectively our results suggest that BRAF-kinase is not an essential gene in RTECs and vemurafenib nephrotoxicity is BRAF-independent. Though it is important to note that AKI is a multifaceted disease^2^ that results from a complex interplay between epithelial^31,50,51^, immune^33^, and endothelial^52,53^ cells. Thus, notwithstanding our studies with RTECs, further work is required to discern the role of immune and endothelial cells in vemurafenib nephrotoxicity.

We propose that vemurafenib inhibits ferrochelatase, which likely contributes to RTEC cell-death and AKI. Ferrochelatase is the terminal enzyme of the heme biosynthesis pathway^45^. FECH is a mitochondrial protein that catalyzes the insertion of the reduced form of iron (Fe^2+^) into protoporphyrin. In human, mutations associated with reduced FECH activity are associated with erythropoetic protoporphyria (EPP) that is characterized by cutaneous photosensitivity and progressive liver damage^45^. Proteomic profiling of leukemia cells previously revealed that vemurafenib but not dabrafenib is a potent FECH inhibitor^43,44^. It was thus speculated that FECH inhibition might contribute to photosensitivity and skin toxicities associated with vemurafenib treatment. Our in vivo studies show that vemurafenib treatment results in a progressive decline in FECH activity and development of AKI two weeks post treatment initiation. Importantly, RNAi mediated knockdown or FECH loss-of-function mutations, cause a striking acceleration in the development and onset of vemurafenib-associated AKI (3-7 days versus 15 days). Furthermore, FECH overexpression in cultured RTECs provided significant protection from vemurafenib-induced cell-death.

In contrast with targeted anti-angiogenic therapeutics^54^ where the entire class of drugs cause hypertension due to ‘on-target’ effects, not all BRAF inhibitors seem to cause AKI^22^. This is supported by the observation that patients treated with vemurafenib have a higher incidence of AKI, with a recent retrospective study suggesting that nearly 60% patient develop AKI within the first three months of treatment^22,25^. We propose that this might be due to the unique property of vemurafenib to inhibit FECH. In support, BRAF gene deletion in RTECs did not result in renal impairment in vivo. While it is currently remains unknown if BRAF is essential in other renal cell types, it is interesting that BRAF inhibition has been suggested as a therapeutic option to prevent podocyte injury^55,56^. As of yet, it is difficult to predict which patients are more likely to develop vemurafenib nephrotoxicity. It would be interesting to examine if genetic polymorphisms or physiological conditions associated with altered renal FECH activity play a contributing role.

Our study raises several questions that warrant future investigations. While we show that vemurafenib mediated FECH inhibition contributes to RTEC cell-death, the underlying mechanisms remain unknown. Altered iron homeostasis is a known feature of AKI^57–59^; however, the role of FECH in RTEC function and AKI have not been previously explored. It is conceivable that vemurafenib-mediated FECH inhibition might result in accumulation of by-products of heme biosynthesis^44^, which might cause direct toxicity to RTECs through increased oxidative stress. Furthermore, the reduction in heme biosynthesis could also result in mitochondrial dysfunction and RTEC cell-death. Additionally, while genetic FECH knockout triggers RTEC cell-death, it is unlikely that FECH inhibition in RTECs is solely responsible for vemurafenib nephrotoxicity. FECH-dependent and independent mechanisms in RTECs and other renal cells including endothelial and immune cells might play a role in this toxicity.

Due to the availability of BRAF inhibitors such as dabrafenib and the moderate nature of vemurafenib nephrotoxicity, currently there is no dire need to develop therapeutic strategies to prevent vemurafenib-associated AKI. However, several approved, experimental, and investigational kinase inhibitors have recently been demonstrated to inhibit FECH activity, through mechanisms that remain elusive^44^. For instance, an ALK (anaplastic lymphoma kinase) inhibitor alcetinib can inhibit FECH^43^ and is linked with renal dysfunction^60^, although the causality remains to be investigated. Since, FECH inhibition is not limited to vemurafenib, but has emerged as a relatively common feature of several protein kinase inhibitors, future genetic and pharmacological studies are required to understand the role of this enzyme in renal function and AKI.

In summary, our work suggests that vemurafenib-associated AKI is an off target toxicity that can be partly attributed to FECH inhibition in RTECs. By identifying the underlying mechanisms, our study reveals that drugs with FECH-inhibiting ability might cause RTEC dysfunction and nephrotoxicity.

## Methods

### Animal Experiments

Mice were housed and handled in accordance with the Institutional Animal Care and Use Committee of the Ohio State University and the University of Tennessee Health Science Center. Details regarding animal procedures are provided in Supplementary methods.

### Cell culture

BUMPT, HK-2, and primary murine RTECs were cultured according to previously described methods^61,62^. Further details of experimental procedures are provided in Supplementary methods.

### Statistical Analysis

Data are presented as mean with SD. P < 0.05 was considered statistically significant. Further details are provided in supplementary methods.

## Acknowledgements

We thank Dr. Yogesh Scindia for critical reading of the manuscript. We thank Dr. Shuiying Hu for assistance with pharmacokinetic analysis. This work was supported by funds from the OSU Cancer Center (to N.S.P.) and NIDDK Grant R01 DK-117183 (to A.B.). Y.B. was supported by postdoctoral fellowships from the Pelotonia foundation and the American Heart Association.

## Author Contributions

Y.B., J.Y.K., L.A.J., M.G., J.A.S, V.S., K.M.H, S.D.B., A.S., A.B., and N.S.P acquired and analyzed most of the data. Y.B., V.S., J.S., K.D.J and N.S.P conceived the study and performed data analysis. N.S.P approved the final version of the manuscript.

