## Supplementary Material for "Nephrotoxicity of the BRAF-kinase inhibitor Vemurafenib is driven by off-target Ferrochelatase inhibition"

1. **Supplementary Figures (1-10)**
2. **Supplementary Methods**
3. **Supplementary References**

**Suppl. Figure 1**

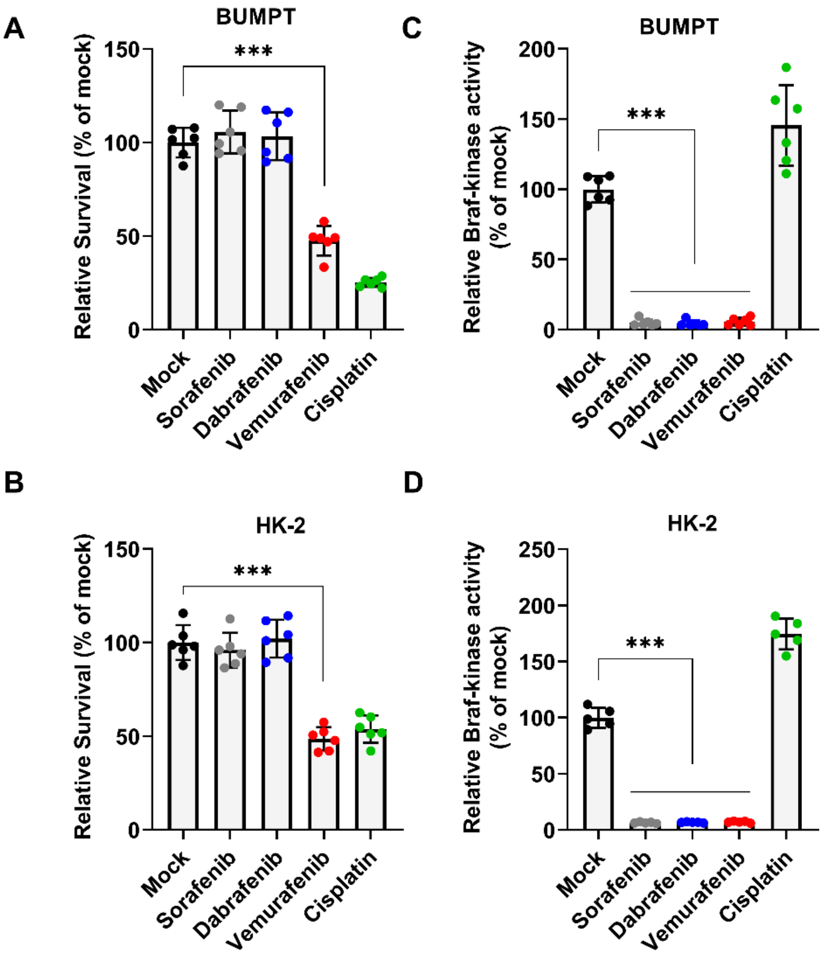


**Suppl. Figure 1: Braf-inhibition and *Vemurafenib induced RTEC cell death.*** BUMPT and HK-2 cells were treated with vehicle or indicated drugs at 50 µM concentration for 48 hours, followed by assessment of (**A-B**) cellular viability by MTT assays and (**C-D**) in vitro Braf kinase assay. For in vitro kinase assays, Braf was immunoprecipitated from whole cell lysates and the kinase activity was normalized to the amount of immunoprecipitated protein. The data indicates that while sorafenib, dabrafenib, and vemurafenib can inhibit Braf kinase activity, only vemurafenib induces cell death in BUMPT and HK-2 cells. In all the graphs (n=5-6 biologically independent samples), experimental values are presented as mean ± s.d. The height of error bar = 1 s.d. and $p < 0.05$ was indicated as statistically significant. One-way ANOVA followed by Dunnett's was carried out and statistical significance is indicated by *$p < 0.05$, **$p < 0.01$, ***$p < 0.001$.

**Suppl. Figure 2**

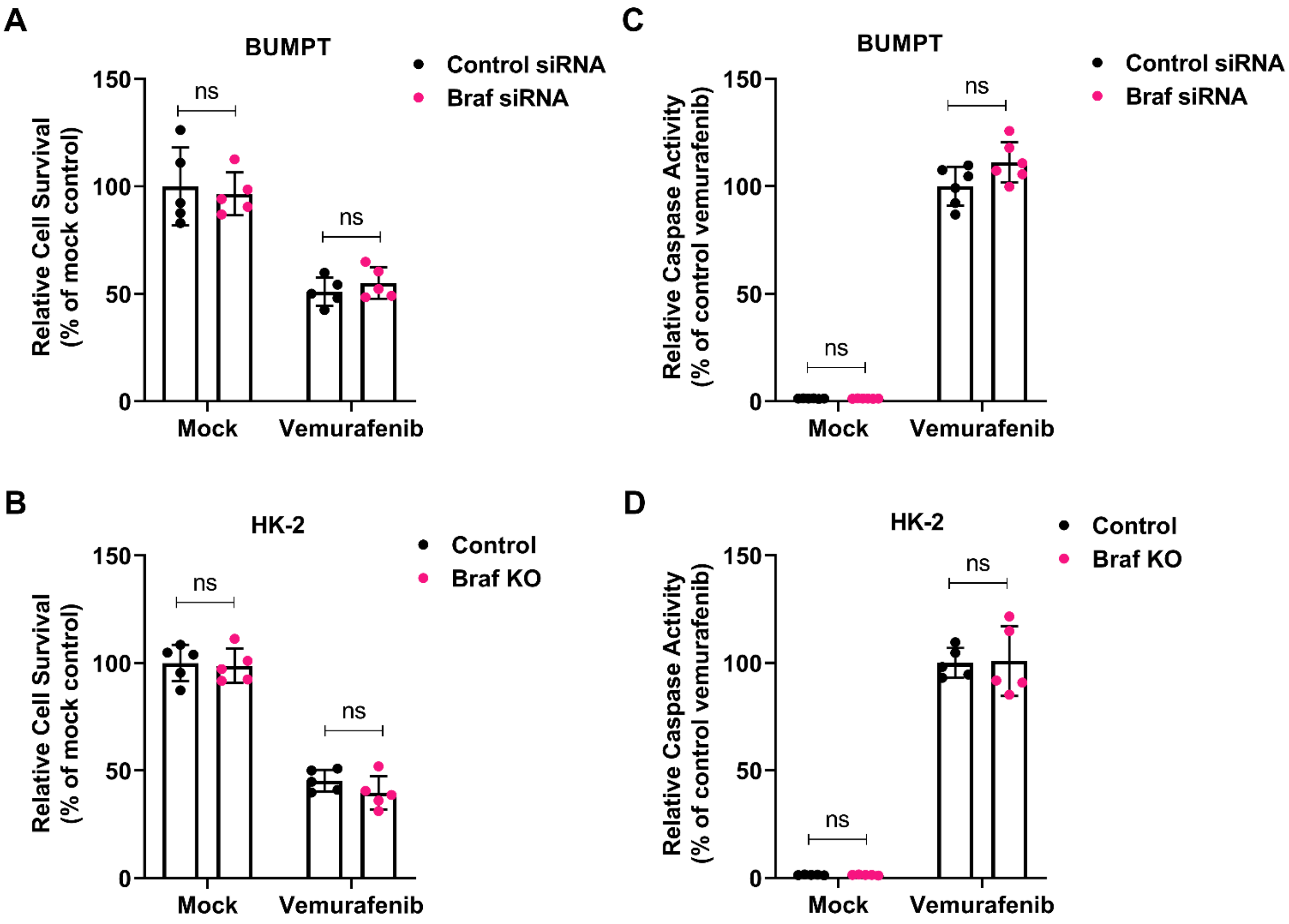


**Suppl. Figure 2:** ***Vemurafenib induced RTEC cell death is Braf independent.*** (**A-B**) RNAi mediated Braf knockdown in BUMPT cells did not influence vemurafenib associated cell death (50 µM for 48 hours) as assessed by MTT based viability assay and caspase activity measurements. (**C-D**) CRISPR/Cas9 mediated Braf knockout in HK-2 cells did not influence vemurafenib associated cell death (50 µM for 48 hours) as evaluated by MTT based viability assay and caspase activity measurements. In all the graphs (n=5 biologically independent samples), experimental values are presented as mean ± s.d. The height of error bar = 1 s.d. and $p < 0.05$ was indicated as statistically significant. One-way ANOVA followed by Dunnett's was carried out and statistical significance is indicated by *$p < 0.05$, **$p < 0.01$, ***$p < 0.001$.

**Suppl. Figure 3**

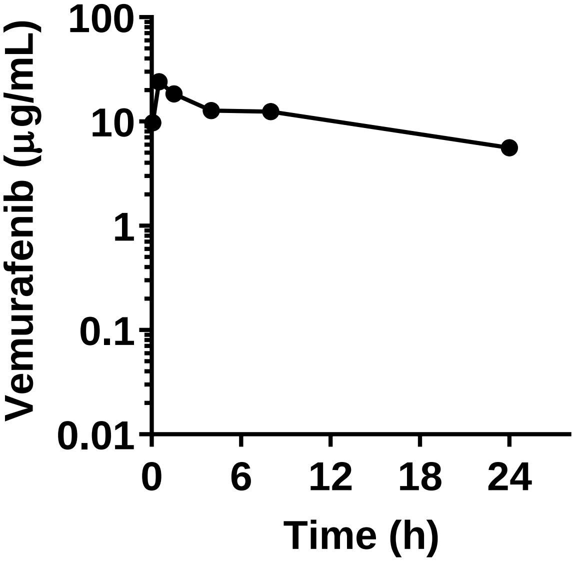


**Suppl. Figure 3:** ***Vemurafenib pharmacokinetic analysis.*** Age-matched, 8-12 weeks old male C57BL/6J mice were injected with 20 mg/kg vemurafenib followed by pharmacokinetic analysis of vemurafenib levels in the plasma.

**Suppl. Figure 4**

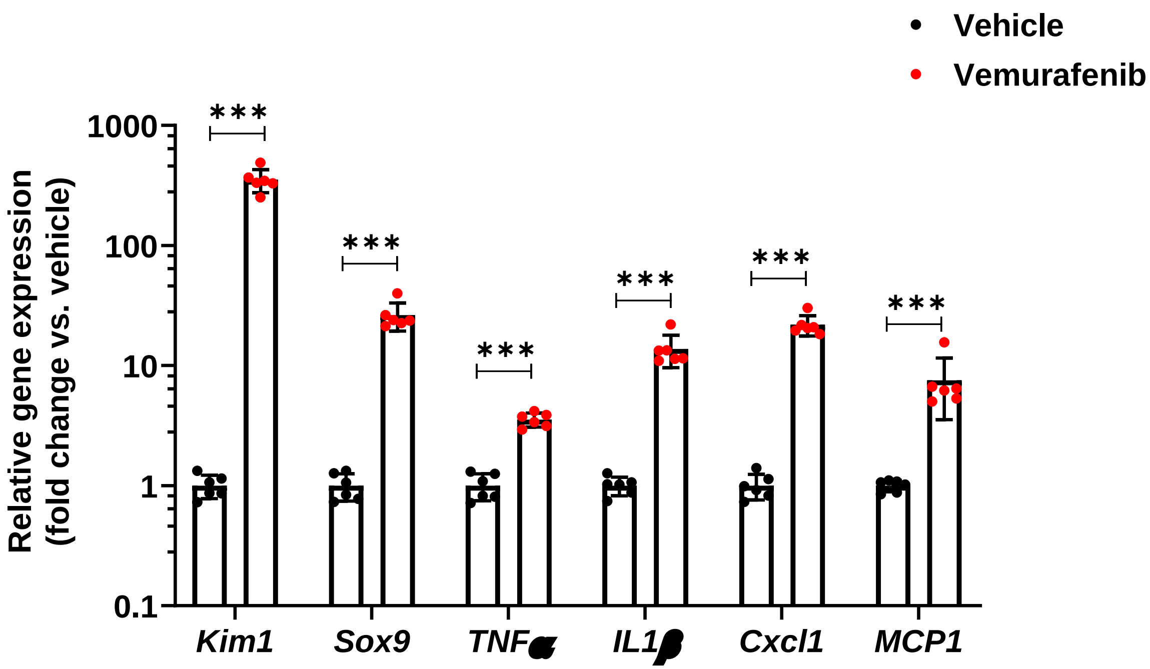


**Suppl. Figure 4: Renal gene expression analysis during *vemurafenib nephrotoxicity.*** Age-matched, 8-12 weeks old male C57BL/6J mice were treated with either vehicle or 20 mg/kg vemurafenib (p.o, b.i.d.) for 15 days followed by gene expression analysis of indicated genes in renal cortical tissues. Injury, inflammation, and repair related genes were upregulated during vemurafenib nephrotoxicity. In all the bar graphs (n=6 biologically independent samples), experimental values are presented as mean ± s.d. The height of error bar = 1 s.d. and $p < 0.05$ was indicated as statistically significant. Student's t-test was carried out and statistical significance is indicated by *$p < 0.05$, **$p < 0.01$, ***$p < 0.001$.

**Suppl. Figure 5**

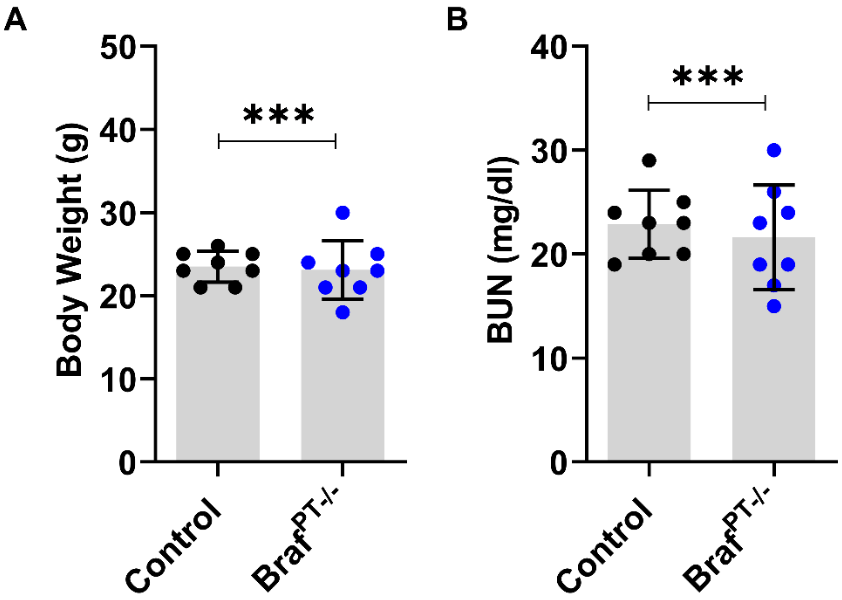


**Suppl. Figure 5:** ***Effect of Braf gene knockout on renal function***. Littermate control and Braf conditional knockout mice (indicated by $Braf^{PT-/-}$) of 8-12 weeks age had similar (**A**) Body weight and (**B**) Blood urea nitrogen levels, indicating that RTEC-specific Braf gene deletion does not influence renal function under normal baseline conditions. In all the bar graphs (n=8 biologically independent samples), experimental values are presented as mean ± s.d. The height of error bar = 1 s.d. and $p < 0.05$ was indicated as statistically significant. Student's t-test was carried out and statistical significance is indicated by *$p < 0.05$, **$p < 0.01$, ***$p < 0.001$.

**Suppl. Figure 6**

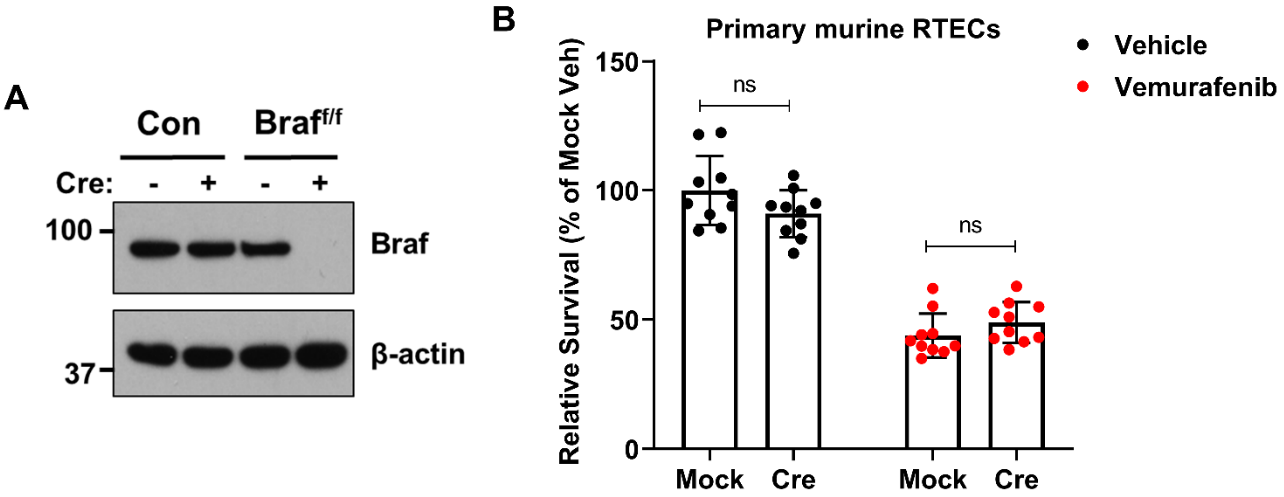


**Suppl. Figure 6: *In vitro Braf gene deletion does not influence vemurafenib associated RTEC cell death*.** Primary renal tubular cells were cultured from control and Braf-floxed mice. One week later, lentiviral transductions (Cre) were carried out to delete Braf gene. (**A**) Immunoblot analysis confirmed Braf deletion. Blots are representative of two independent experiments. **(B)** Primary renal tubular cells from Braf-floxed mice with or without Cre transduction were treated with 50 μM vemurafenib, followed by cell viability assessment at 48 hours using trypan blue staining. In all the graphs (n=10 biologically independent samples), experimental values are presented as mean ± s.d. The height of error bar = 1 s.d. and $p < 0.05$ was indicated as statistically significant. One-way ANOVA followed by Dunnett's was carried out and statistical significance is indicated by *$p < 0.05$, **$p < 0.01$, ***$p < 0.001$.

**Suppl. Figure 7**

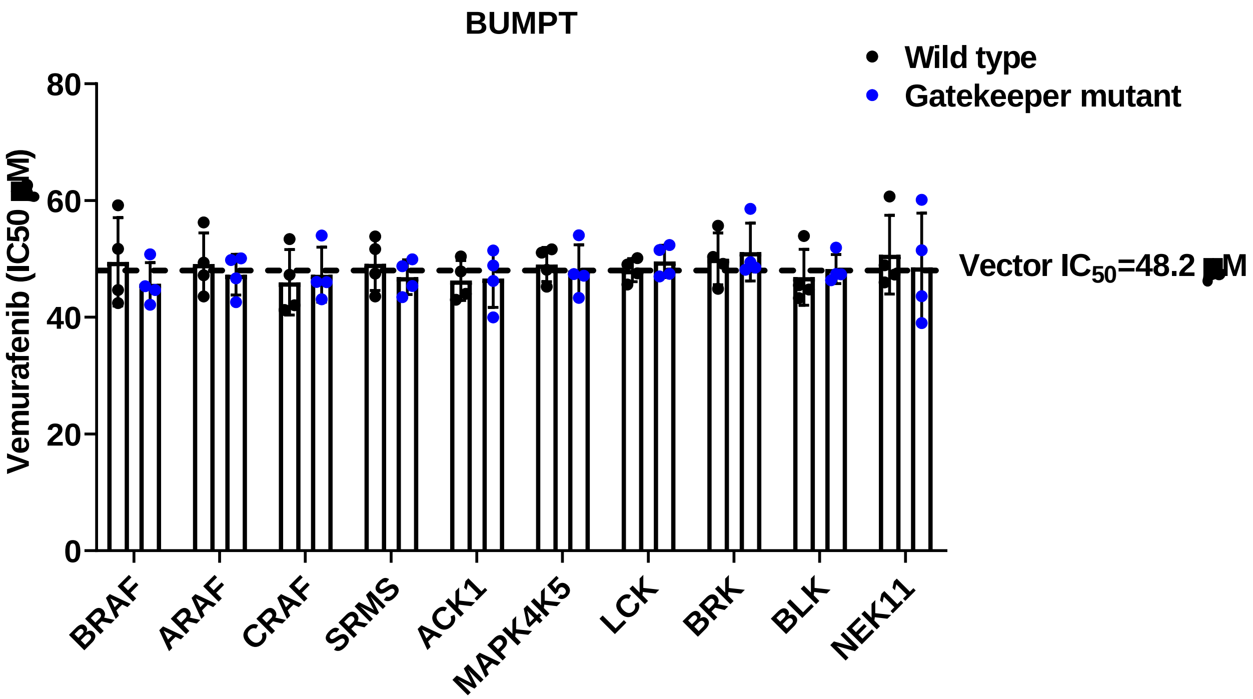


**Suppl. Figure 7:** ***Chemical genetics approach to evaluate the role of kinase inhibition in vemurafenib-mediated RTEC cell death***. BUMPT cells were transiently transfected with empty vector, wild type or gatekeeper mutants of indicated kinases. One day after transfection, the cells were treated with 0-100 μM vemurafenib, followed by cell viability assessment at 48 hours using trypan blue staining. IC50 (half-maximal inhibitory concentration) was calculated by nonlinear regression analysis. The graph represents data from four independent experiments (n=4 biologically independent samples), experimental values are presented as mean ± s.d. The height of error bar = 1 s.d. and $p < 0.05$ was indicated as statistically significant. One-way ANOVA followed by Dunnett's was carried out and statistical significance is indicated by *$p < 0.05$, **$p < 0.01$, ***$p < 0.001$. The results indicate that none of the tested kinases are involved in vemurafenib-associated RTEC cell death.

**Suppl. Figure 8**

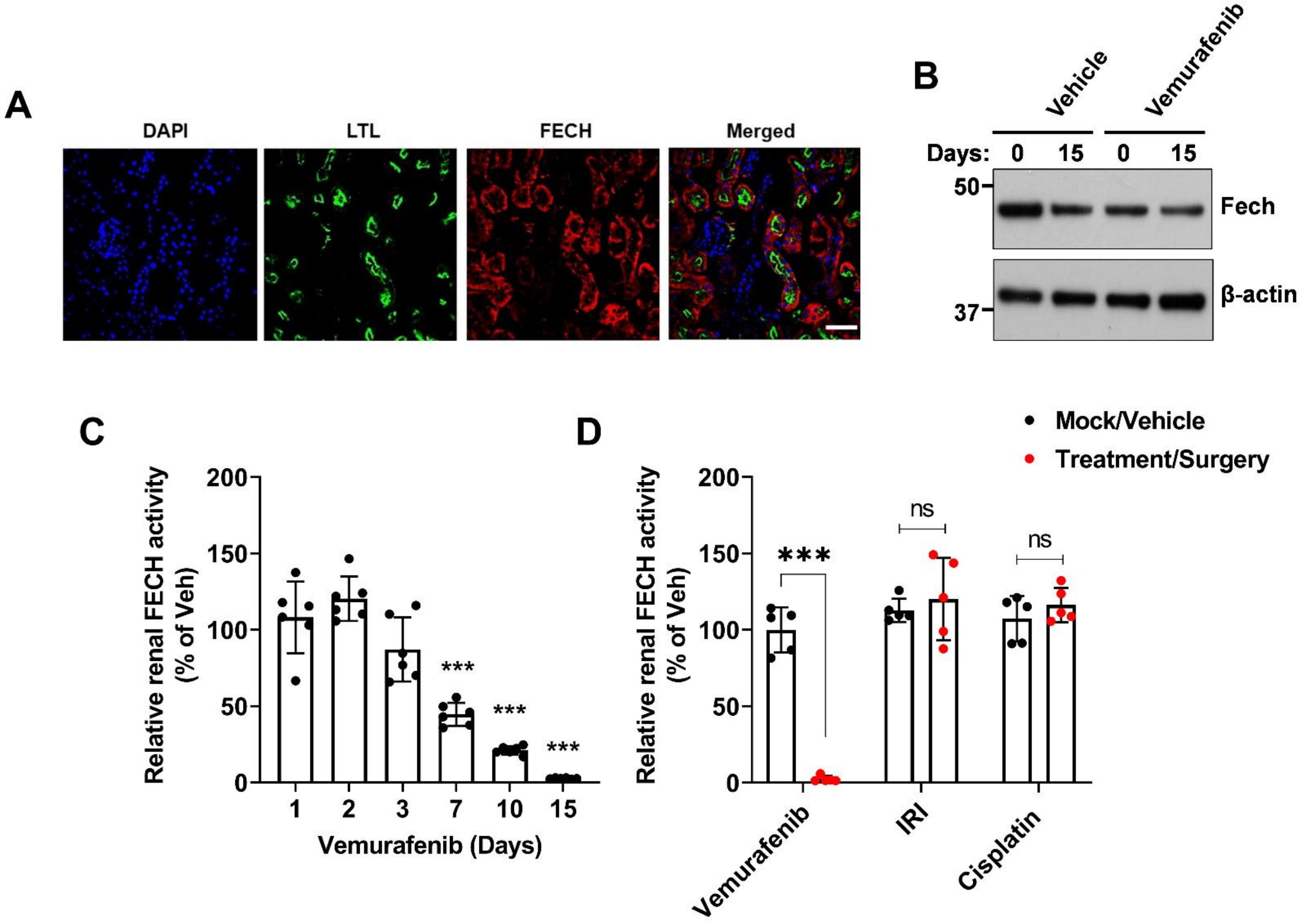


**Suppl. Figure 8:** ***Vemurafenib associated AKI is associated with renal ferrochelatase inhibition.*** (**A**) Immunofluorescence staining of renal tissues showed high FECH staining in RTECs (LTL positive). Age-matched, 8-12 weeks old male C57BL/6J mice were treated with either vehicle or 20 mg/kg vemurafenib (p.o, b.i.d.) for 15 days. (**B**) Immunoblot analysis of renal FECH expression did not show a significant difference in vehicle and vemurafenib groups. (**C**) Renal FECH assay showed a progressive decline in FECH activity in the vemurafenib treated mice. (**D**) Renal tissues from 8-12 weeks old male C57BL/6J mice treated with 20 mg/kg b.i.d. vemurafenib (15 days), 30 mg/kg cisplatin (day 3) or bilateral renal ischemia (30 minutes followed by 24 hour reperfusion) were used to evaluate renal FECH activity. The results show that reduced in FECH activity during vemurafenib-associated AKI is not an indirect effect of kidney injury. In all the bar graphs (n=5-6 biologically independent samples), experimental values are presented as mean ± s.d. The height of error bar = 1 s.d. and $p < 0.05$ was indicated as statistically significant. Student's t-test (A-I) was carried out and statistical significance is indicated by *$p < 0.05$, **$p < 0.01$, ***$p < 0.001$.

**Suppl. Figure 9**

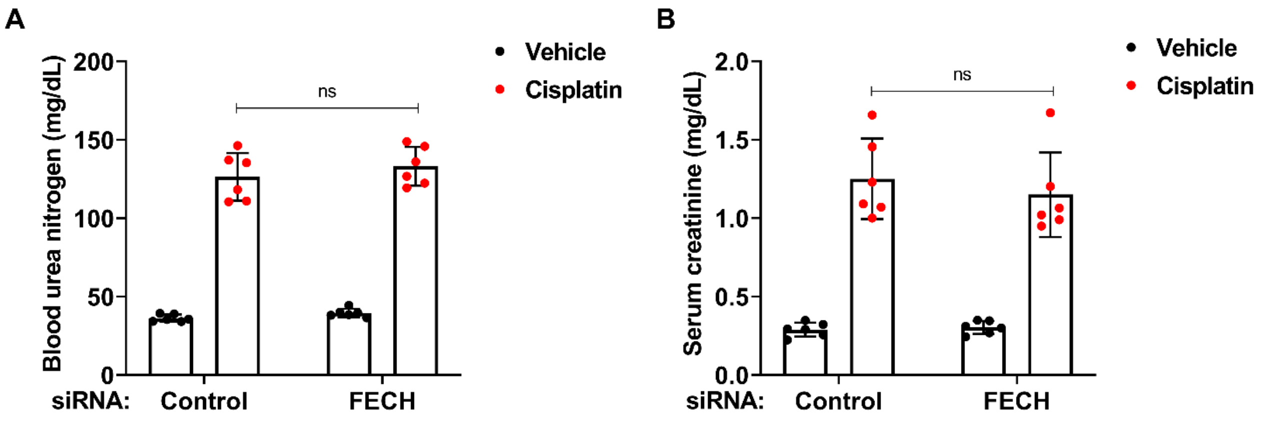


**Suppl. Figure 9:** ***In vivo siRNA mediated FECH knockdown does not influence the severity of cisplatin nephrotoxicity.*** Age-matched male (8-12 weeks) C57BL/6 mice were administered with three once-daily intravenous injections (hydrodynamic) of control (non-specific) or FECH targeting siRNAs (25 µg in 0.5 ml of PBS). One day later mice were treated with either vehicle or 30 mg/kg cisplatin (i.p.), followed by endpoint analysis of renal function at day 3. (**A**) Blood urea nitrogen (**B**), serum creatinine measurement showed that the FECH knockdown did not alter the severity of cisplatin nephrotoxicity. In all the bar graphs (n=6 biologically independent samples), experimental values are presented as mean ± s.d. The height of error bar = 1 s.d. and $p < 0.05$ was indicated as statistically significant. One-way ANOVA followed by Tukey's multiple-comparison test was carried out and statistical significance is indicated by *$p < 0.05$, **$p < 0.01$, ***$p < 0.001$.

**Suppl. Figure 10**

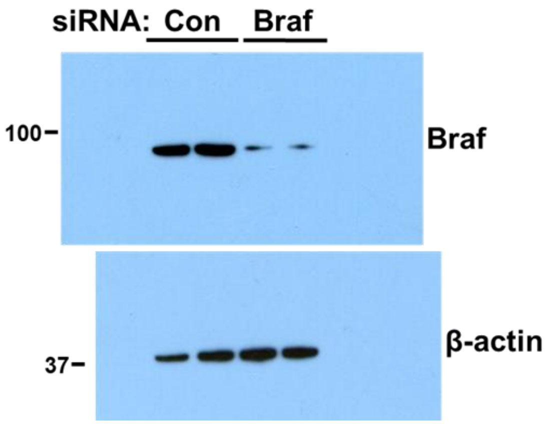


**Figure 1E**

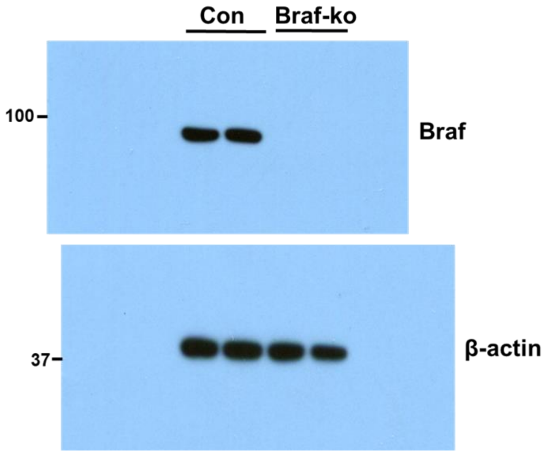


**Figure 1F**

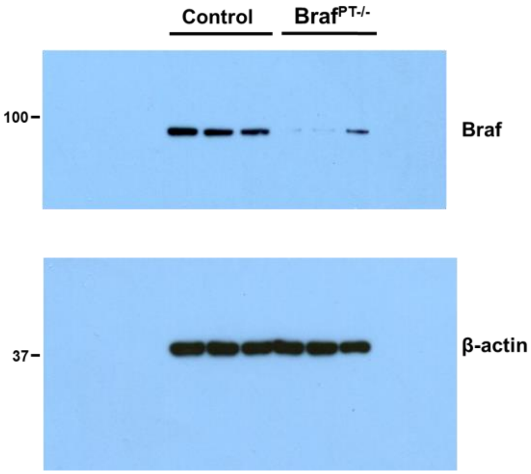


**Figure 3B**

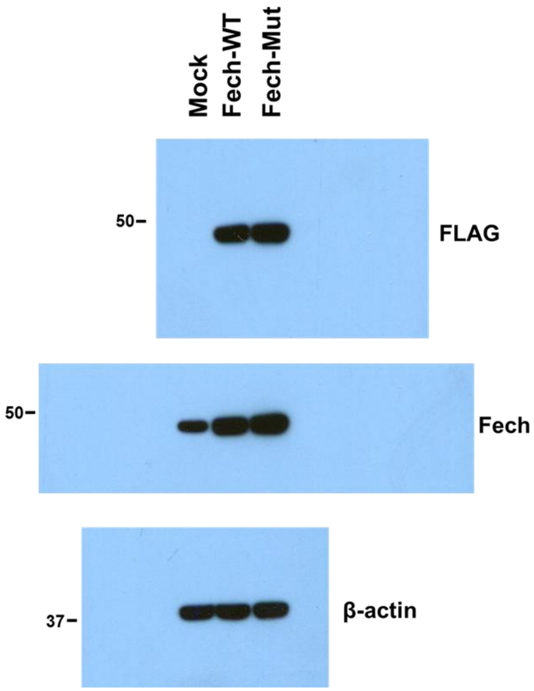


**Figure 4C**

**Suppl. Figure 10:** *Uncropped images of immunoblot data.*

### Suppl. Figure 10 (continued)

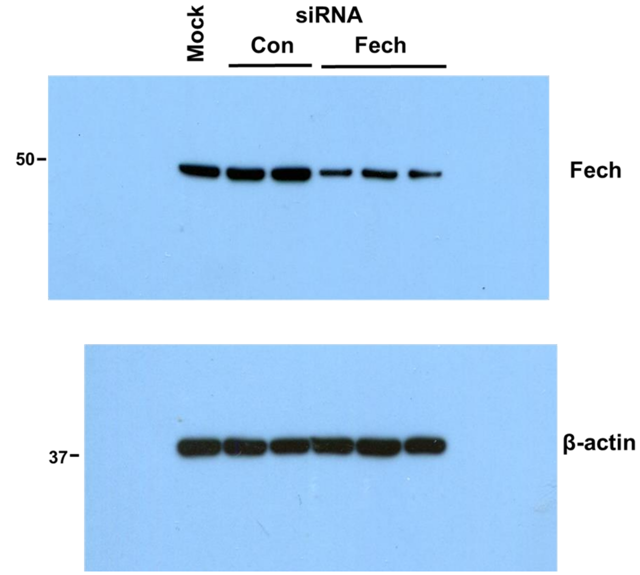


Figure 5A

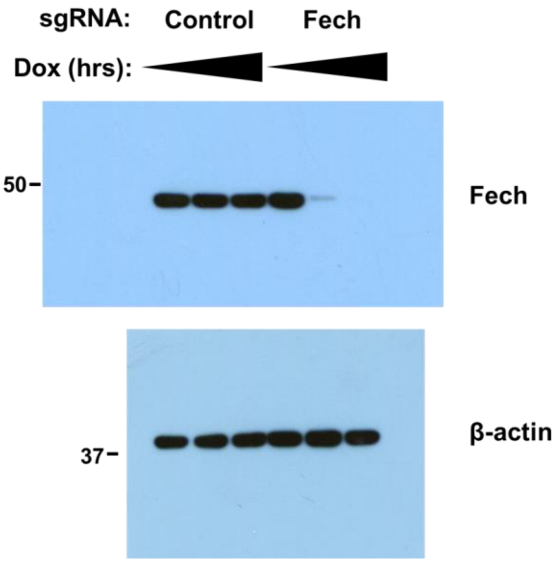


Figure 7A

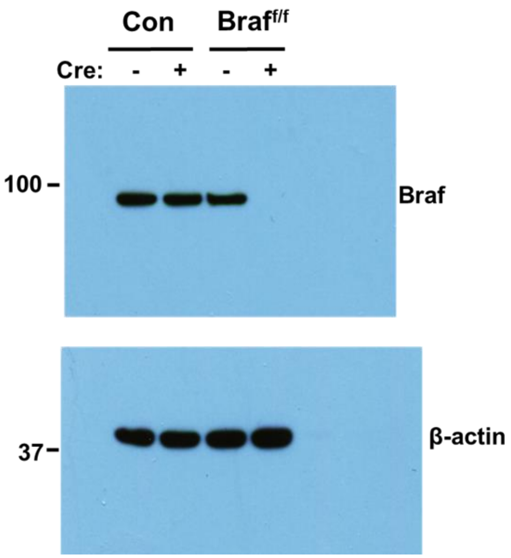


Suppl. Figure 6A

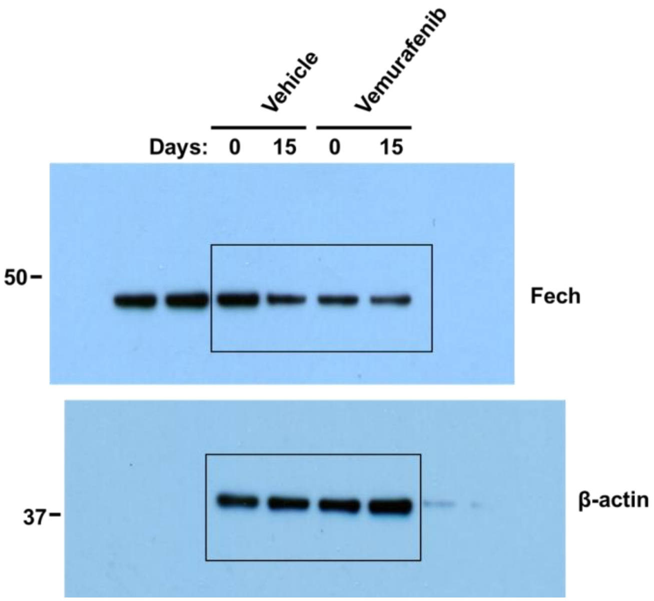


Suppl. Figure 8B

**Suppl. Figure 10:** *Uncropped images of immunoblot data (continued).*

### SUPPLEMENTARY METHODS

***Cell culture and reagents.*** The human renal tubular epithelial cell line, HK-2 cells (CRL-2190) were obtained from American Type Culture Collection (ATCC) and were grown in keratinocyte media (K-SFM) supplemented with 10% fetal bovine serum as described previously. The murine renal tubular epithelial cell line, Boston University mouse proximal tubule cells (BUMPT, clone 306) were generated by Drs. Wilfred Lieberthal and John Schwartz, Boston University School of Medicine, Boston, MA, and were obtained from Dr. Zheng Dong, Augusta University, Augusta, GA). These cells were grown at 37 °C in Dulbecco's modified Eagle's medium with 10% fetal bovine serum as described recently[1]. Cisplatin, vemurafenib and other kinase inhibitors were obtained from Sigma-Aldrich or Selleckchem.

***Kinase gatekeeper mutants.*** The protein kinase plasmids (pCMV6-entry backbone and FLAG tagged) were obtained from Origene. The QuikChange II XL Site-Directed Mutagenesis Kit (Agilent) was utilized to generate mutants, according to methods described in our previous studies[2,3]. The mutagenesis primers for generating the gatekeeper mutations were designed with the help of QuikChange primer design program. Successful mutagenesis was confirmed by DNA sequencing. Lipofectamine LTX (Life Technologies) reagent was used for transient transfection of vector, wild type and gatekeeper mutant plasmids in BUMPT cells for 24 hours, followed by vemurafenib treatment and assessment of cellular viability.

***Primary murine tubular cell culture.*** Primary RTECs were isolated from 6-8 weeks old mice using previously well-established methods[1]. Briefly, after euthanasia, kidneys were removed and renal cortical tissues were minced and digested with 0.75 mg/ml collagenase IV (Thermo-Fisher Scientific). After enzymatic tissue dissociation, cells were centrifuged at 2000 g for 10 min in DMEM/F-12 medium with 32% Percoll (Amersham). Subsequently, the pellets were rinsed with serum-free media and cells were cultured in DMEM/F-12 medium supplemented with 5 μg/ml transferrin, 5 μg/ml insulin, 0.05 μM hydrocortisone, and 50 μM vitamin C on collagen-coated dishes. Approximately one week later on reaching full confluency, the cells were trypsinized and plated at 1 × $10^5$ cells per well in 24-well plates for subsequent experiments. For in vitro Braf gene deletion, primary cells from Braf floxed mice were transduced with high-titer (1 × $10^8$ CFU/ml) LV-CMV-Cre-GFP lentivirus (Kerafast), followed by confirmation by immunoblot analysis at 48 hours. For cell viability and drug treatment experiments, primary RTECs were incubated with 50 μM vemurafenib or vehicle (DMSO) in fresh culture medium for 48 h, followed by viability (trypan blue and MTT) and caspase assays.

***Cell viability and caspase assays.*** For the assessment of cellular viability, we utilized trypan blue staining, MTT, and caspase assays as reported in our previous studies[1]. Transformed RTEC cells lines (BUMPT and HK-2 cells) or primary tubular epithelial cells were plated in 6-well, 24-well, or 96-well plates, followed by treated with appropriate vehicles, cisplatin, vemurafenib, and other kinase inhibitors for 24–48 h. For trypan blue staining, cells from 6-well plates were harvested, followed by trypan blue staining and manual cell counting with a hemocytometer and/or by using Countess Automated Cell Counter (Thermo Fisher). We considered translucent cells as viable and blue-stained cells were considered as non-viable. Finally, cellular viability was estimated by taking the ratio of number of viable cells by the total cell number. In similar experiments where cells were plated in 96-well plates, subsequent to drug treatment, 10 μL of MTT reagent (5 mg/mL MTT in PBS) was added to each well, followed by incubation at 37 °C with 5% CO2 for 4 h. Then, 100 μl of acidified isopropanol (Sigma-Aldrich) was added to each well and absorbance was measured at 590 nm. In certain experiments, IC50 (half-maximal inhibitory concentration) was calculated by nonlinear regression analysis using GraphPad Prism. For measurement of caspase activation as a readout of the extent of cell death, cells grown in 6 well plates were lysed in a buffer containing 1% Triton X-100 to extract cytosolic proteins. 10 μg cell lysates were then added to a caspase assay buffer containing 50 μM DEVD-AFC for 60 min at 37 °C. Fluorescence readings at excitation 360 nm/emission 535 nm was measured, and free AFC standard curve was used to convert the fluorescence reading from the enzymatic reaction into the nM AFC liberated per mg protein per hour as described recently.

***Viral Transduction and CRISPR/Cas9 mediated gene deletion***. Lentiviral transductions were performed using previously described methods[4]. For Cre-mediated gene excision, cultured primary tubular cells were transduced with high-titer (1 × $10^8$ CFU/ml) LV-CMV-Cre-GFP lentivirus (Kerafast), followed by vemurafenib treatment 48 h later. Braf and FECH gene deletion was carried out in HK-2 cells using Lenti-X™ CRISPR/Cas9 System and Lenti-X Tet-On 3G CRISPR-Cas9 System (Takara Bio), followed by isolation of stable cells. Gene knockout was confirmed by DNA sequencing.

***Mice strains and breeding.*** All animals were handled, and procedures were performed in accordance with the animal use protocol approved by the Institutional Animal Care and Use Committee of the Ohio State University and the University of Tennessee Health Science Center. C57BL/6J, Braf floxed, Fech mutant and Ggt1-Cre transgenic mice (stock numbers 000664, 006373, 002662, and 012841 respectively) were obtained from Jackson Laboratories. Braf floxed mice were bred with Ggt1-Cre transgenic mice to generate RTEC-specific knockout mice. The Fech mutant mice have been described and characterized previously[5]. These mice were originally generated through an ENU mutagenesis experiment and were found to harbor a Fech loss-of-function single amino acid substitution mutation (M98K). We bred the heterozygous mutant mice with wild type mice to obtain littermate wild type and heterozygous mice. The pups were ear tagged and genotyped at 3 weeks of age using standard PCR-based methods as described recently[1]. In all the experiments, littermate controls were used.

***Cisplatin and Ischemia-reperfusion associated kidney injury.*** For cisplatin nephrotoxicity experiments, 30 mg/kg cisplatin or vehicle (normal saline) was administered by a single intra peritoneal injection as described previously[6,7]. Subsequently, blood was collected on days 0-2 by submandibular vein bleed and via cardiac puncture after carbon dioxide asphyxiation on day 3. To induce bilateral renal ischemia-reperfusion associated AKI, C57BL/6 mice were administered with ketamine (120 mg/kg, i.p.), xylazine (12 mg/kg, i.p.), and buprenorphine (0.15 mg/kg, s.c.) and placed on a warm pad to maintain the body temperature at 34.5–36°C during surgery. Bilateral flank incision was carried out and the renal vessels on both sides were cross-clamped (26 min ischemia followed by 24 h reperfusion). The Sham mice underwent the same procedure except for vessel clamping as described previously[8,9].

***Vemurafenib nephrotoxicity***. As described previously[10], vemurafenib was dissolved in an aqueous vehicle containing 2% Klucel LF and adjusted to pH 4 with diluted HCl. Vehicle control and vemurafenib were administered orally (0.2 mL per animal, b.i.d., 8 hours apart) at 20 mg/kg dose for 20 days. Subsequently, blood was collected on days 0-19 by submandibular vein bleed for blood urea nitrogen measurement. At endpoint, blood was collected by via cardiac puncture after carbon dioxide asphyxiation and renal tissues were collected for further examination. For hydrodynamic injection, control (non-specific) or FECH targeting siRNAs from Ambion (25 μg in 0.5 ml of PBS; Austin, TX) or 0.5 ml of PBS was injected into the tail vein as described previously[11,12].

***Assessment of renal damage.*** We assessed functional renal impairment and damage through biochemical (blood urea nitrogen and creatinine) and histological analysis (H&E staining). For biochemical analysis, blood urea nitrogen and creatinine measurement were carried out using QuantiChromTM Urea Assay Kit (DIUR-100) and enzymatic assay-based creatinine mesurement (ab65340, Abcam). For histological analysis of renal damage, harvested kidneys were embedded in paraffin and tissue sections (4 μm) were stained with hematoxylin and eosin by previously described methods. For histopathologic scoring, ten consecutive 100x fields per section from at least three mice per group were examined by an investigator in a blinded fashion. The gradation of tissue damage was scored based on the percentage of damaged tubules as described in our previous studies: 0: no damage; 1: <25%; 2: 25–50%; 3: 50–75%; 4: >75%. Tubules that showed dilation, epithelial flattening, cast formation, loss of brush border and nuclei, and denudation of the basement membrane were considered as damaged.

***Vemurafenib pharmacokinetic analysis.*** Pharmacokinetic studies were performed as previously described[13]. Briefly, 8-12 weeks old C57BL/6 mice male mice were administered with 20 mg/kg vemurafenib. Blood samples were collected at various time intervals via submandibular and retro-orbital bleeds, or cardiac puncture into heparin-coated capillaries or tubes. The blood was spun at 12,000 g for 5 minutes to collect plasma. The plasma was stored at -80C until further bioanalytical quantification. The LCMS/MS system consisted of a Vanquish UHPLC system, a TSQ Quantum Ultra triple quadrupole mass spectrometer from Thermo Fisher Scientific, and Thermo Trace Finder General Quan system software (version 3.3). An Accucore Vanquish C18 column (100 × 2.1 mm, dp = 1.5μm, Thermo Fisher Scientific) was protected by a corresponding XBridge®BEH C18, 5-μm guard column. The injection volume of sample was 5.0 μL. The temperature of the autosampler rack was 4°C, and the temperature of the column was maintained at 40°C. Mobile phase A consisted of water with 0.1% (v/v) formic acid and mobile phase B consisted of acetonitrile: methanol (1:3) with 0.1% (v/v) formic acid. The total run time was 5 min. The optimized gradient 1 started at 0-0.5 min with 10% B; 0.5 - 3.0 min, 95% B; 3.0 - 4.0 min, 95% B; 4.0 - 4.1 min, 10% B; 4.1 - 5.0 min, 10% B with a flow rate of 0.4 mL/min. The MS assay setting with the positive voltage applied to the ESI capillary was set at 3500 V, and the capillary temperature was 342°C with a vaporizer temperature of 358°C. Argon was used as the collision gas at a pressure of 1.5 mTorr. Precursor molecular ions and product ions were recorded for confirmation and detection of vemurafenib (490.118 > 254.929; >99% purity, Sigma-Aldrich Woburn, MA), using palbociclib as an internal standard (448.268 > 380.111; >99% purity, Alsachim, North York, ON, Canada). Results from assay validation studies involving quality control samples analyzed over several days revealed that the within-day precision and between-day precision ranged 2.88% - 14.5%, with an average accuracy ranging 105% - 114%. The lower limit of quantification was 5 ng/mL.

***qPCR analysis.*** Quantitative polymerase chain reaction (qPCR) was performed for gene expression analysis as described in our previous work[14]. Briefly, one µg RNA from RTECs or renal cortical tissues was reversed transcribed using RevertAid First Strand cDNA Synthesis Kit (Thermo-Fisher Scientific) and qRT-PCR was run using the QuantStudio 7 Flex Real-Time PCR System (Thermo-Fisher Scientific) using SYBR green master mix and gene-specific primers (Sigma). β-actin was used as the internal control and gene expression levels were determined by the comparative CT ($\Delta\Delta^{CT}$) method.

***Immunoblot analysis.*** We prepared whole cell lysates from renal tissues and *in vitro* cultured RTECs using a modified RIPA buffer (20 mM Tris-HCl (pH 7.5), 150 mM NaCl, 1 mM Na2EDTA, 1 mM EGTA, 1% NP-40, 2.5 mM sodium pyrophosphate, 1 mM beta-glycerophosphate, protease, and phosphatase inhibitors) supplemented with 1% SDS. Following protein estimation, 20-75 µg protein per sample was loaded on Invitrogen Bis-tris gradient midi-gels, followed by transfer to PVDF membranes, incubation with primary and secondary antibodies and signal detection by ECL reagent (Cell Signaling). Primary antibodies used for immunoblot analysis were from Santa Cruz Biotech: Fech (377377) and β-actin (47778) and ECM Bioscience: Braf (RP2011) and were used at 1:1,000 dilution. Secondary anti-rabbit and anti-mouse HRP-conjugated antibodies were from Jackson Immuno-research and were used at 1:2,000 dilutions. Uncropped images of immunoblots are provided in **Supplementary Figure 10**. Using previously described methods[15], densitometric analysis was carried out with Image J software, and the signals of target protein was normalized to β-actin signal of the same sample.

***Braf kinase assay.*** In vitro cultured RTECs or murine renal tissues were lysed with a buffer containing 150 mM NaCl, 1 mM EDTA, 1 mM EGTA, 1% (vol/vol) Triton X-100, 2.5 mM sodium pyrophosphate, 1 mM β-glycerol phosphate, 0.2% (wt/vol) dodecyl β-D-maltoside, and 20 mM Tris (pH 7.5) and supplemented with protease and phosphate inhibitors. These cellular and tissue lysates were then subjected to Braf immunoprecipitation as described in our previous studies. To this end, 1000 µg of protein lysate was incubated with 2.5 µg of IgG (control) or anti-Braf antibody at 4 $^{o}$C overnight, followed by addition of 50 µl of agarose protein A/G beads for 4 hours. Bead-bound immunoprecipitates were then collected by centrifugation and washed with lysis buffer four times. Finally, the beads were added to a protein kinase reaction buffer containing 20 µM ATP and myelin basic protein (Millipore) as substrate and incubated at 30 °C for 30 min. Using recently described methods[1] employing the ADP-Glo™ Kinase Assay (Promega) kit, we measured the Braf kinase activity. This assay is a luminescent detection method that provides a measure of relative kinase activity by quantifying the amount of ADP produced during a kinase reaction. To quantify the level of inmmunoprecipiated Braf protein in each sample, immunoblot analysis was performed after the termination of kinase reaction. Relative kinase activity was subsequently calculated by normalizing the kinase activity (luminescence) to the amount of immunoprecipitated Braf protein. Undetectable activity in the renal tissues of $Braf^{PT-/-}$ mice confirmed the specificity of the immunoprecipitation-based kinase assay.

***Ferrochelatase assay.*** FECH activity was measured by enzymatic formation of zinc-protoporphyrin IX (Zn-PpIX) using a previously described method[16]. Briefly, cell lysates from cultured cells or renal cortical tissues were incubated with 200 µM PpIX (Sigma) in an 200 µl assay buffer (0.1 M Tris-HCl, 1 mM palmitic acid and 0.3% v/v Tween 20, pH 8.0), followed by addition of 50 µL of 2 mM zinc acetate solution. The reaction mixture was incubated at 37°C followed by addition of 500 µL ice-cold termination buffer (1 mM ethylenediaminetetraaceticacid (EDTA) in 30:70 DMSO/methanol). Susequently, the reaction mixture was centrifuged at 14,000 g for 10 min and the Zn-PpIX in the supernatant was measured with a synergy fluorescence plate reader using a 405 nm excitation/590 nm emission filter. For each sample, a heat-inactivated group was included as a negative control. The specificity of the assay was confirmed using Fech mutant tissues.

***Statistical Analysis.*** In the current study, data in all the graphs are presented as mean with s.d. We used GraphPad Prism software for statistical analysis. $p<0.05$ was considered as statistically significant. To evaluate statistical significance between two groups, two-tailed unpaired Student's t test was performed. For comparisons among three or more groups, one-way ANOVA followed by Tukey's or Dunnett's multiple-comparison test was performed. All experiments were repeated atleast three times and no sample outliers were excluded.
